# Origins, History and Molecular Characterization of Creole Cane

**DOI:** 10.1101/2020.07.29.226852

**Authors:** Dyfed Lloyd Evans, Shailesh Vinay Joshi

## Abstract

Since it was first introduced to Europe in 711 CE and planted in the Americas in 1506, a single type of cane dominated sugar production for 1100 years, until it was finally ousted by Tahitian cane c. 1790. This cane became known as ‘CreoleâĂŹ and is present in the ancestry of many sugarcane hybrids, even today. Whether there was only a single variety of Creole cane or multiple varieties has been a matter of debate for decades. Creole cane remains relevant today, as a Creole cane from Jamaica is the currently chosen lecotype for Saccharum officinarum. In this study we identify 18 historical images of Creole cane, many not previously published. We employ image analyses to characterize the internodes and demonstrate evidence for only a single type of Creole in the new world. Chloroplasts and 45s ribosomal RNA sequences from the cultivar BH10/12 (known to have a Creole female parent) were determined that Java ribbon cane is the historical New World sugarcane known as Creole. We demonstrate that Creole cane is an hybrid and not a single species. Thus *S. officinarum* has no type specimen. We also sequence a ribbon cane (also known as Guinguam) that appeared in the Caribbean between 1790 and 1810 and demonstrate that this cane was a Sinense type from Java that links back to the work of Rumphinus (1660s).

## 1 Introduction

For over 1000 years, there was a single type of sugarcane known to the Western world. So ubiquitous was this sugarcane type that it did not even have a name. Only later, during the 1800s when new sugarcane types came from Polynesia and Java was this cane retrospectively given a name ‘Creole’ (or variants thereof, depending on the local language). For convenience, however, we will be using the name Creole for this type of sugarcane throughout this article.

Creole cane was responsible cane sugar becoming a commonplace commodity, rather than an expensive pharmaceutical compound. It also led to large-scale deforestation, the plantation system, slavery and the rise of rum as the spirit of choice in the 18th and 19th centuries. To understand these processes we need to go back to the origins of sugarcane, how it arrived in Europe and how it made its way to the New World.

Today, there can be little doubt that the sugar industry began in India. It is where sugarcane technology (cane cutting knives, roller mills, animal powered mills, juice concentration and sugar crystallization) were first developed. The Sanskrit word ‘sakara’ had attained its new meaning of ‘granular sugar’ by 500 BCE (Galloway 2005) and the Jatakas (folk tale collections written between 600–400 BCE) describe sugarcane cultivation, the use of presses and boiling the juice to concentrate it (Cowell and Chalmers 2017). In the Arthassatra (c. 324–300 BCE) we have the first unequivocal reference to granular sugar being manufactured. The same text also mentions five distinct varieties of sugar (Shamastry 1967). At about the same date (325 BCE) we have an account from Alexander the GreatâĂŹs navarch and admiral, Nearchus (reported in Arrian’s Indica) of ‘a reed in India brings forth honey without the help of bees’ (Arrian 2013). It was almost a millennium later that sugarcane technology reaches China. Between 626-649 CE Indian envoys in Tang China taught sugarcane methods after Emperor Taizong made his interest in sugar known (Sen 2003).

The Achaemenid conquest of the Indus valley (c. 535–323 BCE) brought sugarcane to Persia and Persia became on of the main sources of sugar for Europe (Sen 1999). The Moslem conquest of Persia (651 CE) brought sugarcane into the Arab world and it quickly became a mainstay of Arab cropping systems (Akram 1975). Extension of the Umayyad Khalifate and conquest of Mediterranean states (c. 650–711 CE) brought sugarcane to Syria, Palestine, Egypt, Sicily, Malta, Morocco and the Iberian peninsula (al-Andalus) (Galloway 1977). The combination of irrigation, citrus, sugarcane and other staple crops into a single agricultural cropping system brought about what has been described as the ‘First Agricultural Revolution’ in the Muslim world (Watson 1981).

The Reconquest of al-Andalus (modern Andalucia, in Spain),a period extending from the Battle of Toulouse in 721 to the fall of the Emirate of Granada in 1492, effectively expunged Muslim rule from Europe (OâĂŹCallaghan 2003). Sugarcane culture was effectively lost in Europe (apart from in Portugal) duting this time. However, in parallel with the final stages of the re-conquest the First Crusade (instigated by Byzantine Emperor, Alexios I Komnenos who requested support in his conflict with the Seljuk led Turks) and which continued until 1099 (Asbridge 2004) brought northern Europeans in contact with Arab cropping systems, particularly in the regions of modern Syria. According to orthodoxy, returning crusaders settled in the Mediterranean and they brought sugarcane with them to the now predominantly Christian Mediterranean countries of Cyprus, Malta and Sicily (Deerr 1929).

The island of Madeira, which lies between the Spanish Canary Islands, Morocco and Portugal in the Atlantic ocean was first discovered by Portugal in 1419. By 1420 the Portuguese had begun to colonize these uninhabited islands under the patronage of Prince Henry the Navigator (Greenfield 1977). In 1425 the first sugarcane was introduced into Madeira from Sicily by order of the Infante D. Henrique (Greenfield 1977; Rebora 1968). Sugarcane growing in Madeira required terracing, drainage ditches and considerable manual labour. It was not long before slave labour initially from the Canaries, and then increasingly from Africa was introduced to run the cane fields. This was the inception of the plantation system that was transported along with sugarcane to the Americas (Greenfield 1979). From Madeira the Portuguese, in response to increasing European demand for sugar exported the sugarcane plantation system to the islands of the Azores, the Cape Verdes and SÃčo Tom. These became the models for the eventual establishment of sugarcane plantations in the New World.

In parallel with the Portuguese in Madeira, the Spanish began colonization of the Canary islands. However, unlike Madeira the Canaries had a native population, the Guanches. Castillian conquest of the islands began in 1402 with the expedition of the French explorers Jean de Béthencourt and Gadifer de la Salle, nobles and vassals of Henry III of Castile, to Lanzarote. From there, they went on to conquer Fuerteventura (1405) (Mercer 1980). In 1448 Maciot de Béthencourt sold the lordship of Lanzarote to Portugal’s Prince Henry the Navigator and this began the transportation of Canary island natives as slaves to Madeira. However, the Spanish crown did not agree to this and the Portuguese were expelled in 1459. The Spanish gained complete control of the islands by 1496, where the Canaries came under the control of the Kingdom of Castille (Rumeau de Armas 1975). It was only after the conquest of the Canaries that the Castillians imposed single crop cultivation on the island — initially with sugarcane and then wine. The date and source of the introduction of sugarcane to the Canaries is not precisely know. Some sources claim that it was introduced by the Portuguese in 1420, others that it was directly introduced by Andalucian settlers from Spain (Greenfield 1979). However, the industry must have been established by 1484 when the first mill was built in Grand Canaria (Galloway 2005).

Columbus’ ‘discovery’ of the New World in 1492 changed the political (and sugar production) map of the world completely. Columbus had been a sugar merchant and he was intimately aware of the profitability of the crop. Indeed, he transported sugar cane from the Canary Islands to Hispaniola (now the Dominican Republic) on his second voyage in 1493 (Deerr 1949, Purseglove 1979). This initial planting failed, as did several others over the intervening years. However, the first successful planting was in 1506 based on cane brought from the Canary Islands to Hispaniola (San Domingo). According to Tussac (1808) who quotes the Spanish historian Herrera, it was a certain Aquilon, denizen of the La Vega region of Andaluca (one of the main sugar growing regions of Spain) who brought sugarcane from the Canaries to Saint Domingue in 1506. The first New World sugar cane mill began grinding in about 1516 in Hispaniola. Sugar production spread to Cuba, Jamaica, Puerto Rico, and the other Greater Antilles by the end of the 1500’s (Hagelberg 1985)

The origins of sugarcane in Brazil remain mysterious, even today. Though ‘discovered’ by the Portuguese in 1498, the land was actually claimed for the Portuguese Empire on 22 April 1500, with the arrival of the Portuguese fleet commanded by Pedro ÃĄlvares Cabral (Boxer 1969). It is believed that Pedro Capico brought the first sugar cane plants to Brazil from Madeira, though the date of introduction is unknown (Simonsen 2005). However, by 1526 the Lisbon customs house was collecting duty on the sugarcane coming from Brazil (Varnhagen 1981). The Portuguese introduced the plantation system. Initially they attempted to co-opt the natives as slaves, but they proved recalcitrant and the large-scale transportation of Africans began. By 1570 Brazil’s sugar production rivalled that of all the Atlantic islands, combined (Greenfield 1979). Part of the impetus in terms of Brazilian sugar production was the collapse of the Madeiran industry. This was mainly due to deforestation, and without wood sugar cannot be produced (Moore 2009).

From 1630 to 1654 the Dutch seized productive areas of northeast Brazil, capturing the plantations from the Portuguese. The Dutch brought with them painters and naturalists and from this time we begin to see accurate depictions of sugarcane in art (Bethell 1987).

The main thread of the story of sugarcane now moves northwards to the Caribbean. Though sugar plantations had been established throughout the Caribbean by the Spanish, the expulsion of the Dutch from Brazil brought new people and new technologies to the Caribbean. Though claimed for the Spanish crown in the late 15th century, the Portuguese claimed the island between 1532–1536 but abandoned it in 1620. It was never permanently settled by Europeans until 1625 when it became an English colony. Sugarcane was established and by 1643 it was a major sugarcane manufacturing colony. As a wealthy sugar-producing colony it also became an English centre for the African slave trade (Frere 1768).

Following its ‘discovery’ by Christopher Columbus in 1794 Santiago became part of Spanish rule. They introduced sugarcane, the plantation system and slavery to the island. In 1655 England conquered the island, re-naming it Jamaica (Parker 2011). From the Age of Conquest, we move to the Age of Discovery and Jamaica becomes crucial in the modern story of Sugarcane and its scientific study. Accompanying the new Governor of Jamaica to the Caribbean in 1678, Hans Sloane ultimately only spent 15 months in the region. During this time, he noted 800 new species of plants, which he catalogued in Latin in 1698. He later wrote of his visit in two lavishly illustrated folio volumes. With his second volume of 1707 (Sloane 1707) containing an image of sugarcane. Sloane ultimately donated his collection to the British people and the sugarcane inflorescence he collected in Jamaica remains within the Natural History Museum of London’s collection (Reveal et al. 1998).

When he came to describe sugarcane in his Species Plantarum (Linnaeus 1753) Linnaeus, though coining the name Saccharum officinarum unaccountably had a specimen of Miscanthus floridulus labelled as sugarcane (Linnaeus Herbarium specimen 77.2, LINN). Though he did have a specimen of sugarcane, this had no associated inflorescence and could not be used as a type. Moreover, no complete specimen of sugarcane could be found in any of the collections that Linnaeus referenced. However, in his Hortus Cliffordianus (Linnaeus 1738) mentions that ‘he considered Sloane’s illustration of sugarcane to be excellent’. This prompted Reveal et al. (1998) to retrospectively assign Sloane’s specimen as the type specimen for sugarcane. Thus sugarcane has a type specimen and that type is Creole cane.

Exploration of the Pacific now brought new sugarcane types to the attention, first of botanists, then of sugarcane plantation owners. In 1768 French navigator Bougainville brought his ship, the Boudense into port on the island of Mauritius. There he left a sugarcane that subsequently became known as Bourbon or Otaheite. Cossigny in 1782 introduced canes from Java to both Mauritius and Reunion and, at his own expense, he had these canes sent to the French West Indies (LÃľgier 1901). At the end of his second voyage (in the Providence) captain Bligh arrived at the island of St. Vincent with his cargo of breadfruit and Polynesian sugarcane, amongst which was Otaheite (Deerr 1949). Within a decade these new sugarcanes had replaced the original cane in the new world. The new canes were taller, had longer and broader internodes and a different flower morphology. They also had names (often from where they were obtained — for example Otaheite is a Francophone corruption of Tahiti). This resulted in the original cane of the American requiring a name and it was christened Creole (or local variants thereof). Very rapidly Creole cane passed out of common usage, though it was retained locally as a chewing cane due to its thin rind. The new canes had thicker rinds (thus were harder to mill) but contained much more juice and higher sugar concentrations than Creole cane. Deer (1949) equated Creole cane with Jamaican ‘Ribbon Cane’ (a light green striped cane) and with the pale Puri cane of India (this is in contrast to the Roibbon canes of Batavia and Mauritius, as described by Descourtilz (1829).

In the 19th century it was known as a fact that sugarcane was sterile. Though the plant could flower (as Rumphius had demonstrated (Rumpf 1741)) and as is shown in many contemporary drawings (see Results section), however as Napoleon IstâĂŹs botanist has demonstrated, even though Creole cane produced seed (fuzz) it was all sterile (OâĂŹConnell 2012).

However, in 1858, a plantation worker Iran Aeus Harper noticed some seedlings with serrated leaf edges that must have been from sugarcane. This was reported by the plantation owner, James Parris who, the same year, wrote a letter about this to Kew Gardens. The notion of sugarcane fertility as dismissed by Kew gardens as everyone knew that sugarcane was sterile. At the time, the plantation was growing Boubon (Otaheite), Transparent (Light Preanger, but long mis-identified as Cheribon) and Native (Creole) cane (OâĂŹConnell 2012). The genie was out of the bottle, though and the fertility of Polynesian and Batavian (Javanese) canes could not be disputed. The next major figure in the story of sugarcane fertility is John Redman Bovell, a native Bajan, who came from a family of medical practitioners who were allied to the planters by marriage. Despite his humble origins, he worked his way up to the position of the superintendent of the experimental station at Dodds. In 1888 he observed that sugarcane could, indeed, flower and this time, when he wrote to Kew he was listened to and his discovery caused a stir. As the contemporary Argosy (a Demeraran newspaper) stated: âĂŸA door, revealing a limitless vision, is opened to breeders and experimentersâĂŹ.

Bovell continued his research on the genetic diversity of Otaheite. Then he began to cross known males with known females within a lantern to generate offspring of known parentage. By 1924 Bovell had raised 118 669 seedlings. Though most were not successful, it was he who was responsible, in 1910 for the creation of BH10/12. This new cultivar soon came to replace White Transparent (Light Preanger) as the dominant cane of Barbados (OâĂŹConnell 2012). The origins of BH10/12 lie in Bovell’s original Otaheite x Creole crosses and it seems that, unusually for modern hybrids BH10/12 retains its S. officinarum female parentage, however it is likely that in its immediate ancestry lies a Light Preanger male as well as an Otaheite male.

Though Creole cane has almost disappeared from the world, having been ousted initially by the introduction of the Polynesian canes and then having been ravaged by sareh disease (Artschwager and Brandes 1958) the cultivar BH10/12 having a Creole female parent retains its Creole chloroplast and retains the history of its hybrid nature in its 45s ribosomal RNA complement. As a result, it is possible to confirm the identity of a putative Creole cane by sequence comparison with BH10/12. This would also allow the confirmation that Creole cane was an hybrid. Due to some of the uncertainties as to where the original sugarcanes brought by European colonizers to the Americas came from some, such as Sauer (1966) and Daniels and Roach (1987) challenged as to whether there was only a single âĂŸCreoleâĂŹ in the New World, or whether Creole represents multiple genotypes. This is an open question that will be addressed in this article. Even the hybrid/non-hybrid nature of Creole cane has not been resolve. This is an important question, as if Creole cane is an hybrid of S. officinarum and S. spontaneum (as many of the early sugarcanes from India seem to be) then it cannot serve as a type specimen for S. officinarum.

At the time that Bovell was creating his hybrids in Barbados, the POJ (Proefstation Oost Java) was also creating hybrids. Initially they imported *S. barberi* cultivar Chunnee from India and hybridized this with the noble barieties Striped Preanger and Cheribon to yield the seedlings: POJ 33, 36, 105, 139, 213, 234 (Paliatseas 1955). Like Creole cane *S. barberi* cv Chunnee is also an hybrid of *S. officinarum* and *S. spontaneum*. Elucidating the ancestry of Chunnee has significant implications in understanding the origins of modern sugarcane hybrid cultivars.

## 2 Results

Historical Perspective

The question of where the Creole cane of the New World came from is a vexed one. Carefuly reading of the literature indicates that the sugarcane in Sicily was probably not introduced by the Crusaders, but remained the same type as that introduced by the Umayyad Moslems. The writings of AbÅń al-Fadl, a celebrated orthodox jurist in the capital of Qairouan in North Africa (in todayâĂŹs Tunisia), refused to eat a cake made with sugar which came from Sicily because Sicily was then ruled by the Fatamids, a Muslim dynasty originally from Egypt, who he considered apostates (Galloway 1977).

Moreover, when the island was captured by the Normans in the late eleventh century the sugar industry still existed but its fortunes had begun to fluctuate. The Norman chronicler Hugo Falcandus (late 12th century) makes several references to sugar production in Sicily and describes not only the sugarcane plantations, but also molasses cooking and sugar refining. There were sugar cane mills in the twelfth century, and in the thirteenth century King Frederick II restored sugar refineries in Palermo, after having to send for experts in Lebanon, indicating that expertise had disappeared and the mills must have been in ruin (Galloway 1977). The logical conclusion is that the sugarcane of Sicily remained that introduced by the Moslems several centuries earlier. There was not real break in sugar cultivation on the island, though the technology for sugar manufacture may have lapsed.

Despite sources to the contrary, we cannot be exactly certain of when sugar cane was first planted in Madeira and from where it was sourced. Most of the sources claim that it was introduced from Sicily at the instigation of Prince Henry (de Cadamosto 1944; Dias Leite 1947). Some authorities, however, doubt this since the crop already was being raised in Portugal. Still others, like Duarte Leite, believe that the Genoese, already present as merchants in Lisbon and Madeira, introduced the crop on their own from the Mediterranean (Leite, 1941). Regardless of this, the sugarcane grown throughout the European Mediterranean remained the same as that introduced by the Arabs between 711 and 900 CE. These were the same sugarcanes as introduced into Madeira and the Canaries by 1500 CE.

Image Identification In total, 19 historical images of sugarcane were identified, including two previously unidentified European images from the 14th and 15th centuries. The first of these (Figure 1a) is the oldest European image of a sugarcane in flower. It is also the first image to show a sugarcane grower (with a cane knife) and a sugarcane merchant. A full reproduction of all images and their documentary sources are given in Supplementary Document 1. These very earliest images are not diagnostic, as they are drawn stylistically and resemble *Arundo donax* more than they do sugarcane. However, they also follow the style of the very first image of sugarcane known in flower. This comes from the Arabic world and probably slightly pre-dates the European image and shows a single stool of sugarcane with five Sticks. The image derives from a palimpsest deposited in the BibliothÃĺque Nationale de Paris MS arabe 2771. It is a portion of a treatise on agriculture by Ibn al-BatÄĄr bound into the Book of Simples. In both these images, the flower is shown as a dense, ovate, panicle a feature that can be followed as sugarcane enters the New World.

**Fig. 1.**
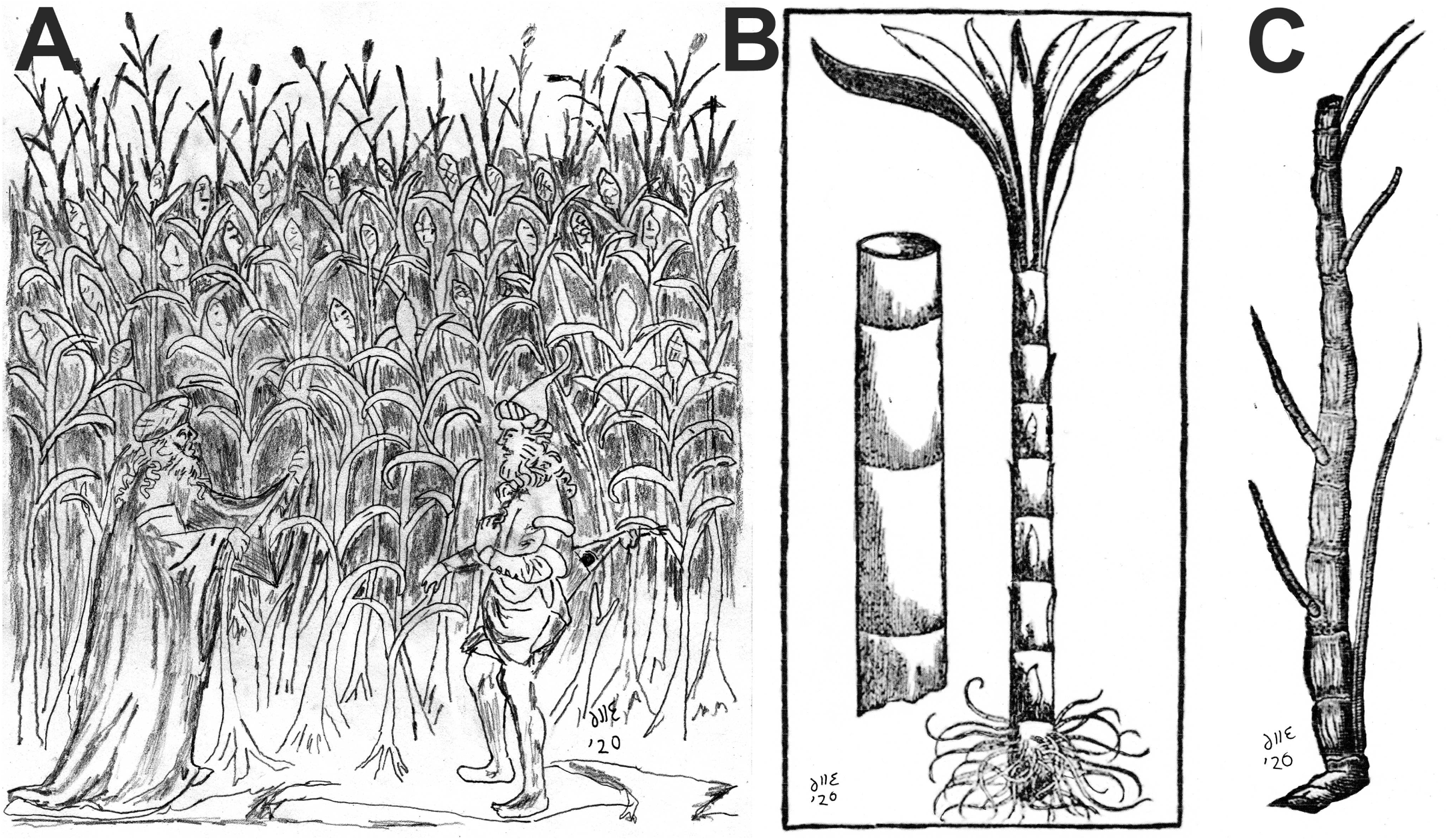
Earliest representations of sugarcane in European art. The first image (A) is the first image of sugarcane and sugarcane in flower in European art. Like most of the earliest representations of sugarcane it was found in *Tacuinum Sanitatis* (Handbook on Health) Folio 92v, which not only shows sugarcane in flower but also depicts the first representation of a cane grower and a cane merchant, with the grower having a cane knife in a pouch on his back. The second image (B) is a wood-cut from MÃijnster (1570)âĂŹs *Cosmographia Universalis*. The final image is another woodcut, this time from DalÃľchamps (1587) *Historia Generalis Plantarum*. All images re-drawn by and copyright D Lloyd Evans (reproduced with permission).

Image Analysis In European art, the first recognisably modern image of sugarcane is found on the page describing Malta in the 1570 edition of Sebastian MÃijnsterâĂŹs Cosmographia Universalis (first published in 1544). The image (Figure 1b) depicts a mature sugarcane stalk, with an accurate image of the roots and the presence of buds on alternating internodes though the leaves are depicted a little stylistically. It has to be remembered that this image was produced before the enlightenment (perspective etc) and it was produced as a woodcut for printing thust it may not be completely botanically accurate. A more accurate depiction is given in DalÃľchampsâĂŹ Historia Generalis Plantarum (1587); this image is also reproduced in M. de LobelâĂŹs 1591 Plantarum seu stirpium icones vol I, where the plant is named Harundo saccharina indica. Though the main image is of two young sugarcane plants growing from a sett there is, inset, a mature sugarcane stalk (Figure 1c). This is an European cane, from ???? and represents the cane brought to Europe by the Umayyads.

Analyzing the images from two European and 12 New World images of Creole, it is apparent (Table 1) that the majority of images show canes with tumescent internodes and relatively broad nodes. Nodes are about 2âĂŞ3 times as long as the internodes are broad. The exact dimensions of the nodes are hard to determine as there are no external references. The exception to this is the oil on canvas painting of âĂŸMulatto Man with Gun and SwordâĂŹ 1641 by Albert Eckhout. The original is currently in the National Museum of Denmark. Daniels and Roach (ref) interpreted this image as possessing slender and cylindrical internodes. However, computational analysis of a very high resolution image of the painting reveals some clearly tumescent internodes. Shading being the key here, which if excluded gives slightly bobbin-shaped internodes. But when shading is included within the internode shape they are clearly tumescent. Comparing stalk width to hand size gives a stalk diameter of about 3.5cm (given a man of standard height and build for the time period).

**Table 1.**
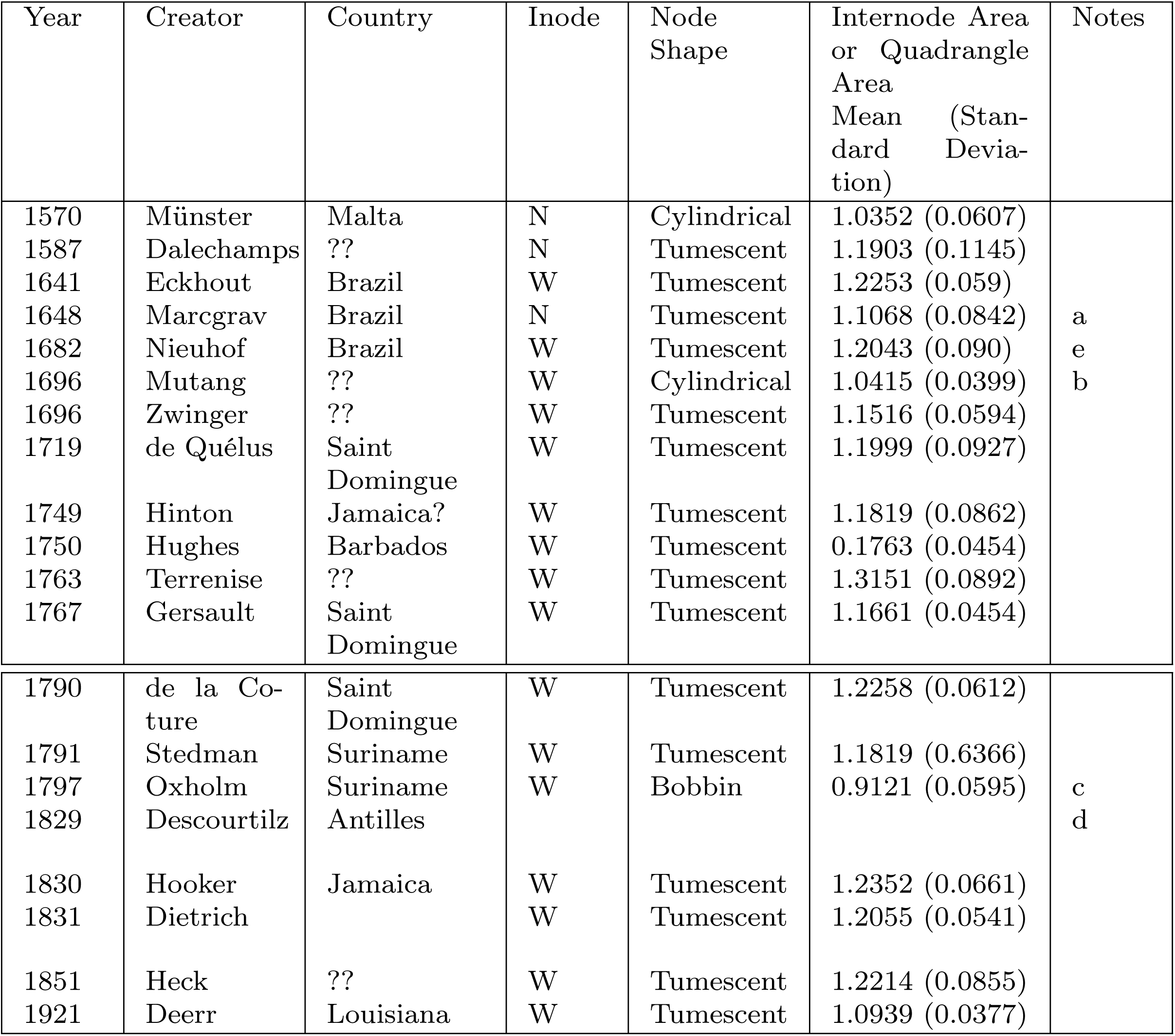
Table giving the internode analysis comparing image analysis to quadrangle area analysis for 17 historical depictions of sugarcane. In the column headings Inode stands for internode shape. Column 5 gives the ratio of the volumes as determined by Icy to the area of the internal or bounding quadrangle. If this measure is >1.1 the internode is tumescent (barrel shaped). Notes: a) though this is a woodcut and to the naked eye the internodes look cylindrical they are actually on the lower end of tumescent. b) The origin and varacity of many of Muntang’s illustrations is in question. His depiction of sugarcane is one of these. The image is included as an exemplar as the internodes are clearly cylindrical. c) this is not Creole cane, but it has bobbin-shaped internodes and is used as an exemplar for the values obtained for this internode shape. d) this is ribbon cane, it’s added to the table to complement our analysis of ribbon (Guingham of Mauritius) cane in this paper. e) Though originally published as an engraving of ‘Plants and trees in Brazil’ in Johan Nieuhof’s Gedenkweerdige Brasiliaense zeeen lant-reize. Amsterdam, 1682 this image actually comes from a pull out re-publishing of the image in Anon (1704) A collection of voyages and travels, some now first printed from original manuscripts. Vol. II. London: Awnsham and John Churchill, Paternoster Row. The double horizontal line marks the period c. 1790 when Polynesian cane arrived in the Caribbean.

Of all the images, only the wood-cuts of Munster and Marcgrav did not support sugarcanes of medium size with tumescent internodes. Due to the nature of wood-cuts, the dimensions of these images cannot be taken as reliable. Thus, from our analyses of the available sugarcane images we find no support for there having been two independent types of Creole in the New World. Moreover, the images of Stednab (1793) presented as âĂŸThe Sugar Cane in its four different stagesâĂŹ from his book âĂŸNarrative, of a five years’ expedition, against the revolted Negroes of SurinamâĂŹ shows how misleading artistic images can be. The mature sugarcane, presented in the centre has apparently cylindrical internodes. These become slightly tumescent in appearance when examined as a high quality scan, but are clearly tumescent in the expanded image of the sugarcane stalk shown in the same image.

One of the major works that seems to have been overlooked by sugarcane scholars over the centuries is the book of de la Coture (1790). Working in Saint Domingue (modern day Haiti) he made a study on the origins, culture and cultivation of sugarcane. This book was recognized by the French Royal Academy of Sciences. In the book he has a magnificent pull-out with a full botanical analysis of sugarcane. The image shows a young sugarcane in flower and a mature sugarcane stalk. These images are shown in Figure 2 and they help explain some of the mysteries surrounding Creole cane. According to de la Coture, the image of sugarcane in flower is from a young specimen (5-6 months old) and it is these young plants that, when given the right conditions do flower. However it is the 15-16 month old plats that are harvested. Figure 2 also compares an earlier image of flowering sugarcane taken from the engraving ‘A Representation of the Sugar-cane and the Art of Making Sugar’ originally published for J. Hinton in the Universal Magazine 1781. The image shows sugarcane in flower, with the same morphology as that of de la Coture. The final image on the top is that of the famous image of Creole cane by Fleischmann (1848). Thus this image of Creole cans is undoubtedly that of an immature plant. The bottom row of Figure 2 shows mature sugarcane plants. First the image of Eckhout 1641, then the image of Piso and Marcgrav (1648) and finally the reference image of de la Coture (of known and verified age).

**Fig. 2.**
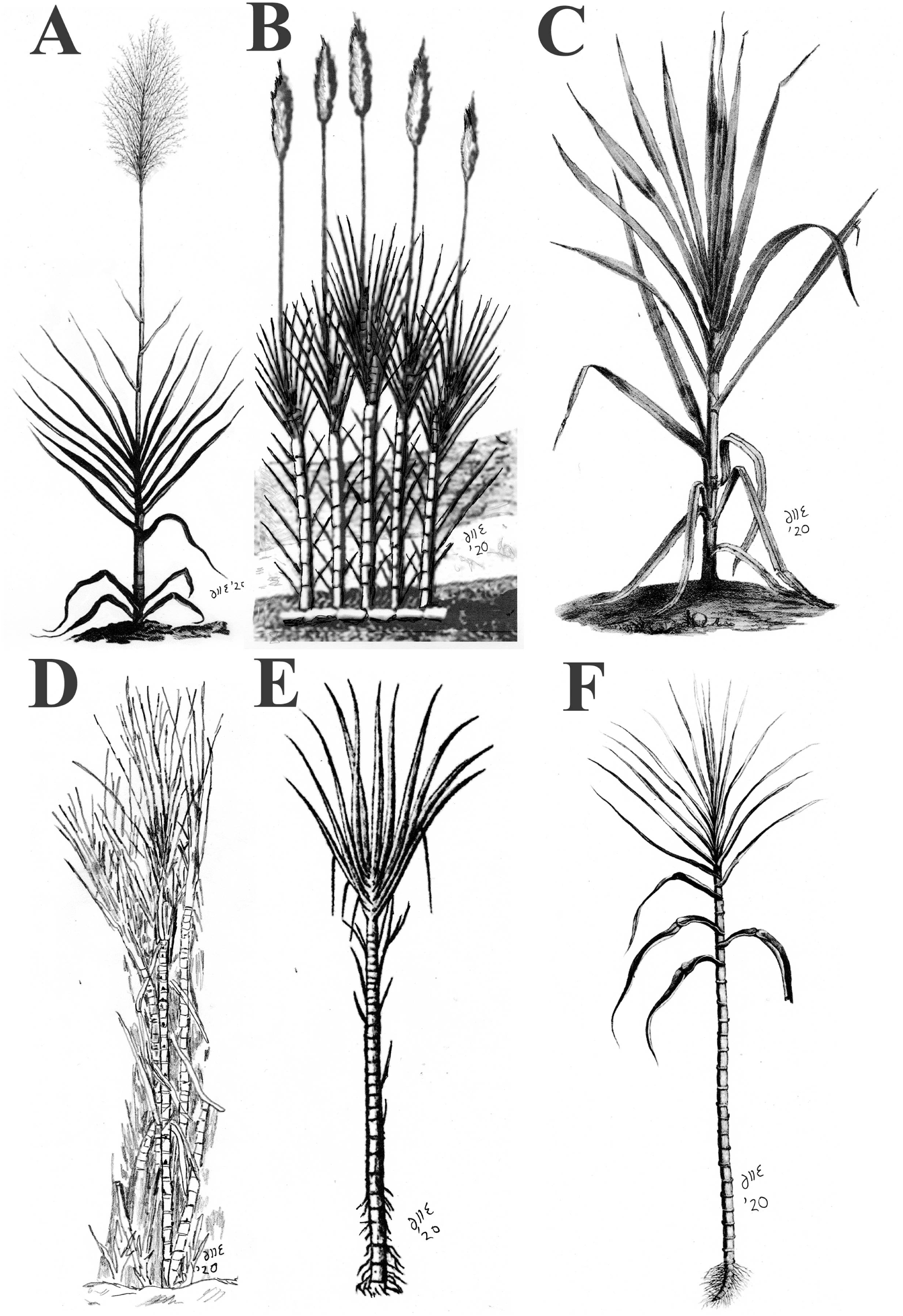
Following de la Coture (1790) it is possible to divide Creole cane images into the two main growth stages of the plant. Top images represent young (4–5 month old) Creole cane in flower. Bottom images represent mature Creole cane (14–16 months of age). Images originate from (A) de la Coture (1790), (B) Hinton (1749) A representation of the sugarcane and the art of making sugar. (C) Fleischmann (1848); (D) Eckhout (1641); (E) Piso & Marcgrav (1648) and (F) de la Coture (1790). All images re-drawn by and copyright D Lloyd Evans (reproduced with permission).

As can be seen there is a significant difference between the immature flowering plants and the mature sugarcane stalks. De la Coture also gives sectional batons from plants of different age and different positions in the plant. We see that the more mature batons have less tumescent internodes than the younger plants. This is entirely consistent with our findings and explains why there has been considerable historical confusion over the images of Creole cane. It should also be noted that Polynesian, Javanese, Mauritian and even Indian canes began arriving in the New world from about the 1780s. Any image before this must have been Creole. Any image after this point needs to be investigated to confirm the identity of the plant being depicted.

On the Biological Nature of Creole Cane. The cultivar BH10/12 along with Jamaican ribbon (Creole) cane and a potential pink-flowered Creole cane from the Bahamas. Complete chloroplast as well as full-length 45s ribosomal RNA cassettes were obtained. BH10/12 was employed as our exemplar as we know from the historical records that it possessed a Creole female parent. Thus its chloroplast sequence should be derived directly from Creole. Employing magnetic bead capture around tRNA genes and MinION sequencing (Lloyd Evans and Hughes 2020) the BH10/12 chloroplast assembled as two single sequences, both of 141 177 bp in length. The differences between the sequences being an inversion of the short single sequence region (Figure 3). Jamaican ribbon cane had an identical topology and differed by 5 single amino acid changes, all in the SSC region of the chloroplast, compatible with SNPs (indeed, the base from BH10/12 was present as the minor allele. The two isoforms of this chloroplast are shown in Figure 3. The potential Creole cane from Barbados was also 141 177 bp long and demonstrated 3 sequence variants as compared with BH10/12.

**Fig. 3.**
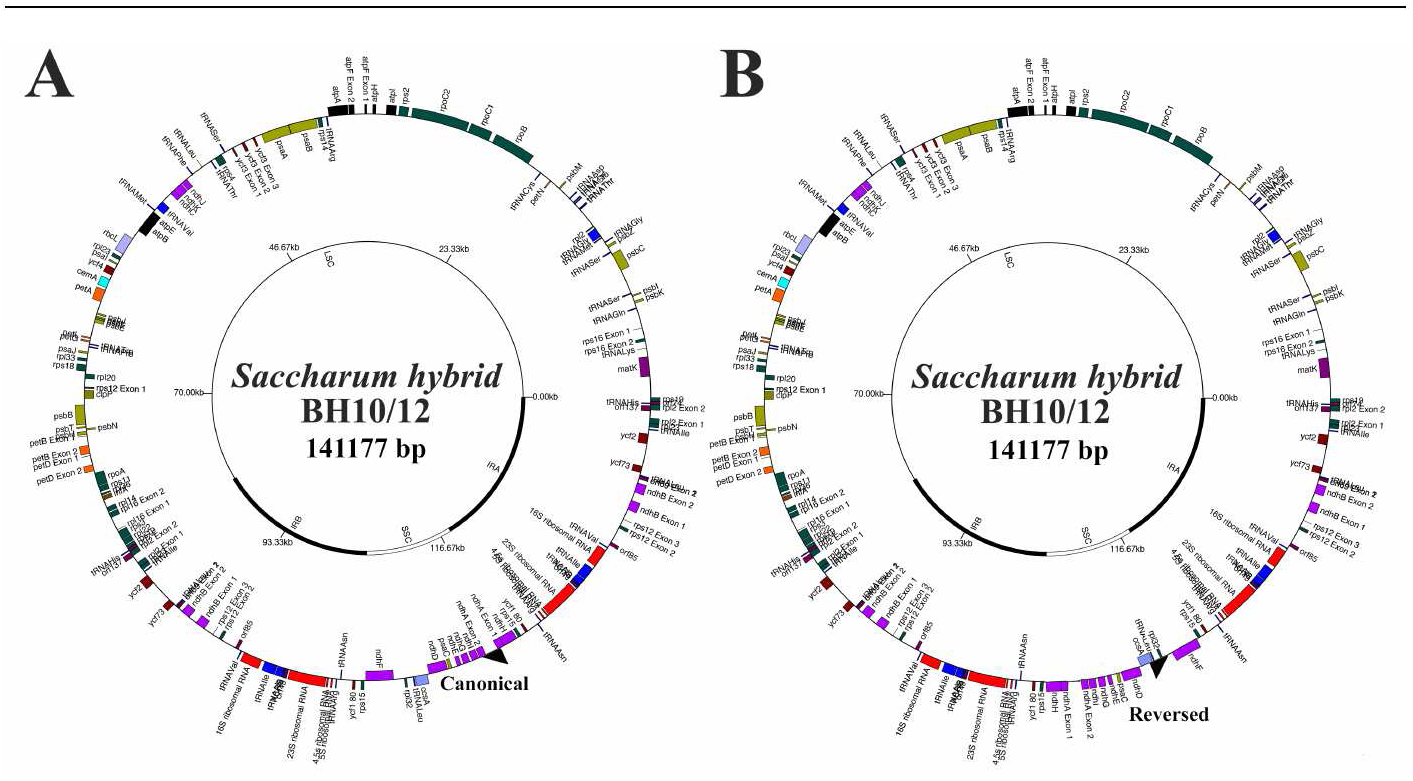
Schematic image of the two chloroplast isoforms of *Saccharum* hybrid BHl0/12. As *Saccharum* hybrid BHl0/12 was employed as our known Creole chloroplast exemplar, a schematic representation of its chloroplast is shown. Genes are marked on the outside. Image (A) shows the standard (canonical) orientation of the chloroplast. However, long read (MinION) assembly also indicates the presence of a second isoform (B) where the small single copy (SSC) region is inverted.

Phylogenetic analysis (Figure 4) demonstrates that all three chloroplast sequences fall within the *Saccharum officinarum* clade, being sister to *Saccharum officinarum* IJ76-921. As in our previous analyses, the *Saccharum officinarum* clade contains *Saccharum robustum, Saccharum barberi* and *Saccharum sinense* as internal members, meaning that none of these three are species in the true sense, they are merely continuations of *S. officinarum. Saccharum cultum* is sister to *S. officinarum* and *S. officinarum* and *S. cultum* are together sisters to *S. spontaneum*. Together these species form *Saccharum sensu stricto* (in the strict sense). *Saccharum s.s.* is sister to a clade formed by African *Miscanthidium* and *Erianthus giganteus* from the Americas (which is the type species for *Erianthus*.

**Fig. 4.**
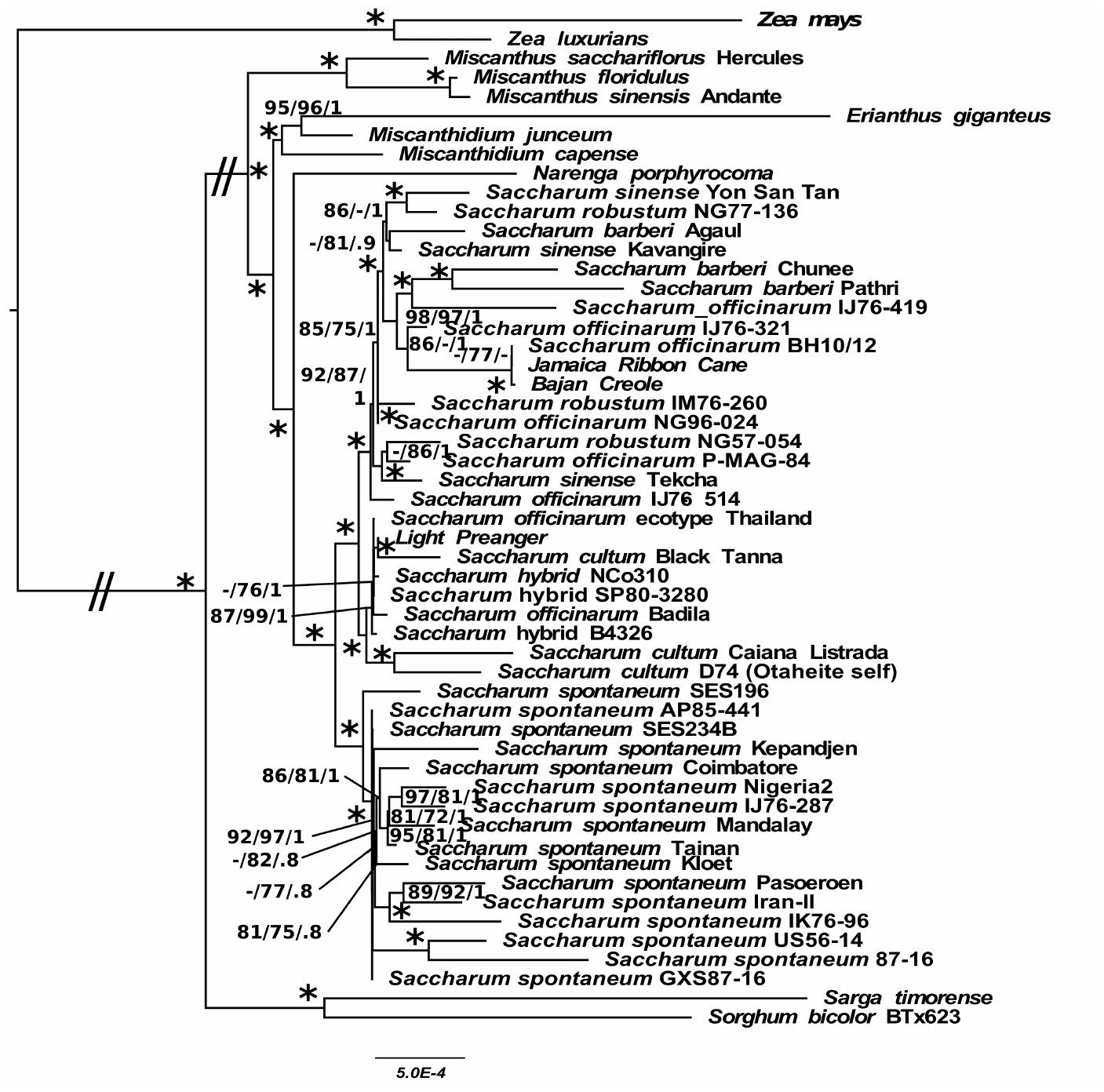
Chloroplast Phy logeny of BHl0/12 and Creole Canes within *Saccharum.* Phylogeny based on whole chloroplast analyses showing the placement of BHlO/12 and the Creole canes within genus Saccharum. Nodes marked* have 100% support across single branch SH-aLRT, non-parametric bootstrap and Bayesian Inference support. Numbers next to nodes represent branch supports in terms of SH-aLRT, non-parametric bootstrap and Bayesian Inference support. A dash (-) shows that branch support is lower than the accepted cutoff (85% for SH-aLRT, 80% for bootstrap and 0.75 for Bayesian Inference). The scale bar on the bottom represents the ex pected number of substitutions per site. Bounding bars on the side delimit the *Saccharum spontaneum, Saccharum officinarum* and *Saccharum cultum* species clades. The double slash (//) marks represent long branches that have been reduced in length by half.

ITS sequence capture and sequencing isolated four ITS isoforms from BH10/12, two from Jamaica ribbon and three from the Bajan Creole. Alignment and phylogenetic analyses of these sequences with other isolated and sequenced ITS regions (Figure 5). All three sugarcane types are hybrids of *S. officinarum* and *S. cultum*. As expected from BovellâĂŹs crossing experiments the additional ITS sequences identified in BH10/12 are compatible with two distinct subtypes of S. cultum. The Bajan Creole also possessed an S. cultum type ITS isotype.

**Fig. 5.**
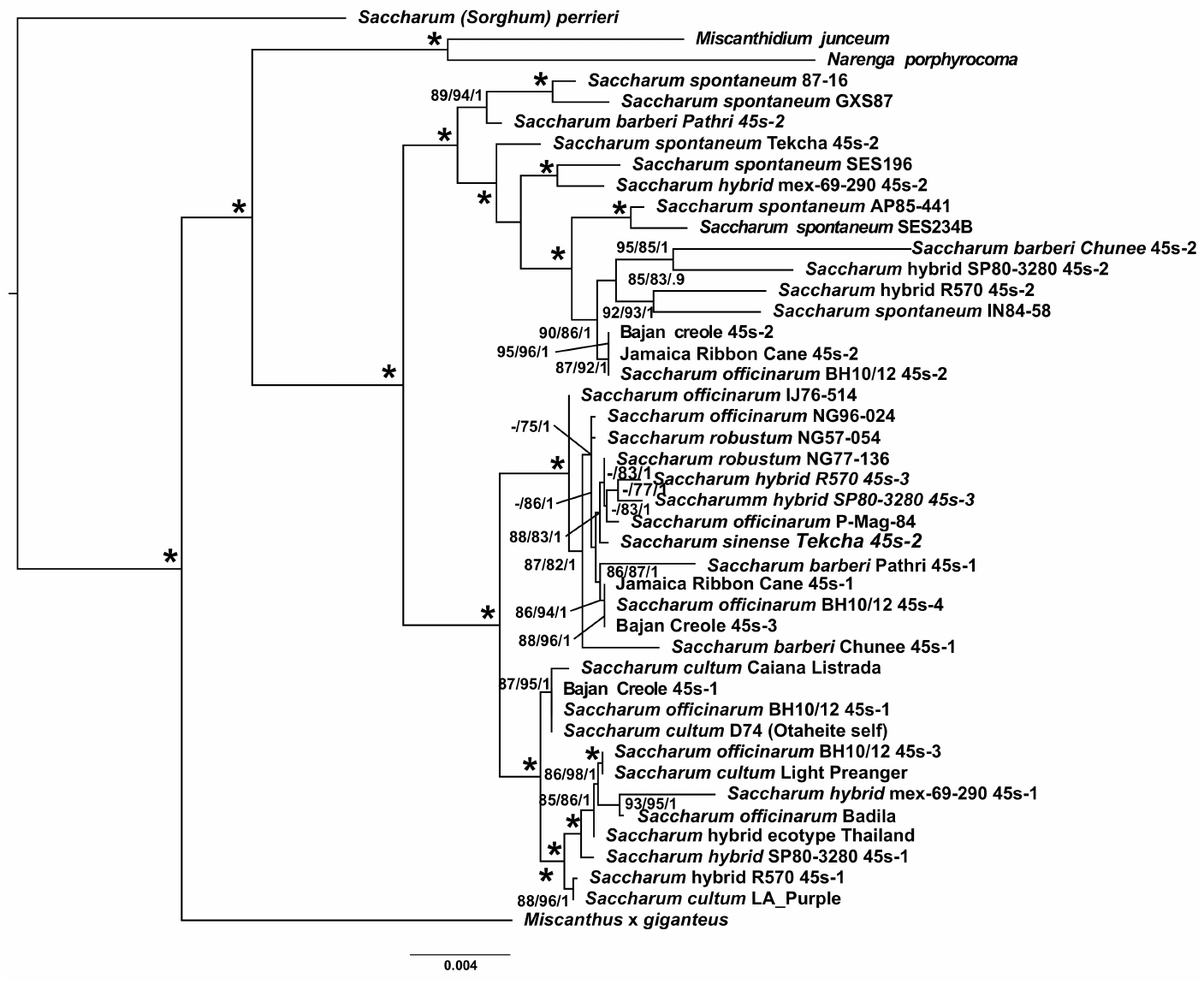
45s Ribosomal RNA Phylogeny of BHl0/12 and Creole Canes in Relation with Saccharum. Phylogeny based on the complete 45s Ribosomal RNA cassette showing the different I TS components found within BHl0/12, Creole cane, âĂŸSinenseâĂŹ and âĂŸBarberiâĂŹ canes as well as mo dern hybrid cultivars in comparison with known reference exemplars. Nodes marked * have 100% support across single branch SH-aLRT, non-parametric boot strap and Bayesian Inference support. Numbers next to nodes represent branch supports in terms of SH-aLRT, non-parametric bootstrap and Bayesian Inference support. A dash (-) shows that branch support is lower than the accepted cutoff (85% for SH-aLRT, 80% for bootstrap and 0.75 for Bayesian Inference). The scale bar on the bottom represents the expected number of substitutions per site. Bounding bars on the side show divisions into *Saccharum spontaneum, Saccharum officinarum* and *Saccharum cultum* species bins.

Intriguingly, the cane image that previous authors (Daniels and Roach 1985) maintained was cylindrical (that of Piso and Marcgrav (1648)) turns out to be tumescent when image analysis is applied. But the image of Deer (1921) turns out to be more cylindrical. There are good psychological reasons for this. Humans are not good at perceiving negative shading in constrast to a clear reference. Thus, to the naked eye the woodcut of Piso and Marcgrav looks as if it has cylindrical internodes. However, in the negavive shading the nodes are clearly pinched in and the internodes are tumescent. The image of Deer is the only one in colour to be analyzed. To the naked eye this image clearly shows tumescent (barrel shaped) internodes. But if the image is redrawn or analyzed computationally the internodes are actually mutch flatter than they appear. The shading gives an illusion of three-dimensionality which is not completely captured in the flat image. Thus, Deerr’s image of Creole cane comes just under the limit of a tumescent internode. Due to the colour shading, however, a scale factor should be applied to this image.

In our analysis, all the historical images of Creole cane had tumescent internodes. The only exceptions being the early European woodcut of sugarcane from Münster with cylindrical internodes and the image of Muntang, which historians (even from his own day) (ref) have considered suspect. Image analysis therefore supports there only having been a single type of sugarcane with medium-thick tumescent internodes grown in the New World before the arrival of Polynesian canes. It was this singular cane that came to be known as ‘Creole’. This is also supported by the historical evidence as all the canes in the New World came from Madeira and the Canary islands and these all appear to have been a single type of sugarcane.

## 3 Discussion

The origin and identity of the first sugarcanes to come to Europe, the selfsame sugar canes that were brought to the new world has long been a matter of debate. These sugarcanes dominated the sugar industry for 300 years (in the New World) and 800 years before that in Europe. This cane was of medium thickness and had a soft rind, making it ideal for crushing methods. However, it had little juice and was relatively poor in sugar (as compared with the canes that ultimately displaced it).

Employing a specialized search bot, we identified the first image of this sugarcane in European art (Figure 1) as well as 12 additional images of good botanical quality that had not been reported previously. Isolation of the cane pictures from the other elements of the images using Photoshop masking, followed by image analysis with ICY (Table 1) revealed that all but two of the images had tumescent internodes (this includes images that had previously been reported as cylindrical). Only two sugarcane images with cylindrical internodes were identified. In the first, the artist may not have seen a sugarcane in the wild. In the second, the sugarcane was almost certainly drawn from nature, but was then transferred to woodcut by a carver who may never have seen sugarcane and the carver’s own prejudices and the ease of carving straight as opposed to curved lines may have influenced the final image. We have also seen from the drawings by de la Coture that the shape of Creole cane internodes do vary with the age of the cane.

We solve a historical mystery (Figure 2), by demonstrating early flowering of sugarcane and maturation at between 5 and 8 months, yielding a stunted cane. However, it the cane does not flower it matures at 15 to 16 months of age.

Sequence (Figure 3) and phylogenetic (Figure 4) analysis on BH10/12 sugarcane which has a known Creole female parent and it was used to confirm that both Jamaica ribbon cane and Bajan Creole also had true Creole female parents. Apart from a few SNPs (9 and, 5 respectively) the chloroplast sequences were identical, demonstrating the same parentage. Jamaica ribbon cane only had S. officinarum and S. spontaneum 45s ribosomal RNA sequences, compatible with what is known for Creole. BH10/12 had two additional ITS sequences, one almost identical to B74 (a known self of Otaheite) and a second compatible with a Javanese S. cutlum (Figure 5). This is compatible with the crossing of Creole with Otaheite and with a Javanese cane (which represents the known parentage of BH10/12). Bajan Creole has S. officinarum and S. spontaneum ITS sequences dominating. However, there is a small but detectable component from a Javanese-type S. cultum sugarcane. The most parsimonious origin of Bajan Creole is that the original Creole cane was crossed with a Javanese cane (as female parent) this was then back-crossed to Bajan Creole (giving a majority Creole inheritance, but with increased disease resistance and the unusual pink colour to the inflorescence).

## 4 Conclusions

The precise nature of Creole sugarcane has been in doubt for many decades as, despite being the main sugarcane type known in Europe from 711 CE to the 1770s it was supplanted first by Polynesian canes and then by Javanese canes. Thus, it had passed into historical obscurity before the molecular tools became available to properly analyze it. However, as this paper has shown, Creole cane formed the foundation on which modern sugarcane breeding was developed and its genetics (or the genetics of related wild hybrids) still lie at the heart of many modern cultivars.

Our extensive image analysis demonstrates categorically (and mathematically) that apart from two outliers all images demonstrate that only a single type of Creole cane was grown in the New World. Computational analyses (Table 1, Figure 2) show that mature canes all have tumescent internodes. Unlike modern hybrids, the work of de la Coture shows that Creole canes flower when the plant is still young… and this may explain the two differing morphologies that some workers have ascribed to Creole cane. The only other mention of sugarcane in early records is in the letters of Alonso de Zuazo, (1466 – March 1539) a Spanish lawyer and colonial judge and governor in New Spain and in Santo Domingo. He writes in a letter to Charles V of Spain that ‘there were excellent plantations of sugarcane, some growing as thick as a man’s wrist. How wonderful it would be if large factories for making sugar could also be built’ (Thomas 1997). Given the context, these words contain a considerable dose of hyperbole and were there for a political reasons. They do not provide any proof for a thick Creole cane in the New World.

Sequence analysis demonstrates that the three identified Creole or Creole derived sugarcanes have identical chloroplast sequences (apart from SNPs) thus they share a single common female ancestor. As reported in the literature BH10/12 is an hybrid with an Otaheite female parent as well as a female parent from Java (compatible with its also having been crossed with Light Preanger). Jamaica ribbon is a classic hybrid of S. officinarum and S. spontaneum, which is compatible with the natural hybrids of India. However, as this is now demonstrated to be an hybrid and this is the same cane (Creole) as that described by Hans Sloane (Sloane 1707), with this image becoming the lecotype for *S. officinarum* (Reveal et al. 1998) this means that a hybrid is the currently selected type specimen for S. officinarum. As a result S. officinarum has not valid associated type. This has been mooted before (ref) but we not demonstrate that this is the case molecularly. The five hundred year old mystery of Creole cane has finally been solved and we have demonstrated that only a single type of Creole cane was present in the New World. Our findings also have implications for the typing of S, officinarum, as the currently chosen type specimen is an hybrid and not a species.

## 5 Methods

### Image and Keyword Search

As more libraries and historical collections digitize their holdings, the number of digitally available historical texts and images are growing dramatically. A search bot was written using the Perl (https://metacpan.org/release/libwww-perl) API. The bot was written to access and protocol sites and to respect robots.. Sites were traversed over a period of 12 months, with a limit per site per day to prevent over-loading the site. The bot searched for matches to a keyword list that was extended over the period of the project. Images, metadata, associated text and were directly downloaded. were converted to text and input into a proprietary natural language processing system (Lloyd Evans and Joshi 2020). The final subset of images and texts were examined manually.

To progress to image analysis, image had to have a verifiable provenance (source and date as a minimum). Images were re-drawn by DLlE for publication and a complete list is given in Supplementary Document 1. Original images were masked in PhotoShop, exported as high quality tiffs and imported into Icy 2.0 (de Chaumont et al. 2012). Using the active contours tool characteristic features such as node heights, node widths, and cross heights were evaluated for each image. Exemplar images from late Middle Age Europe to 1880s Caribbean were selected for further analysis. For the determination of internode areas, the area between the left and right side of the image were approximated by a series of thin horizontal rectangles. The area is then approximated by:

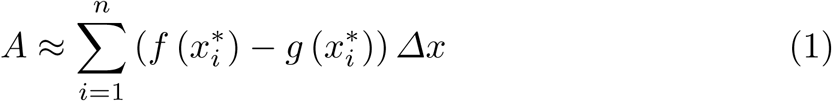

As the limit approaches infinity, the exact area is:

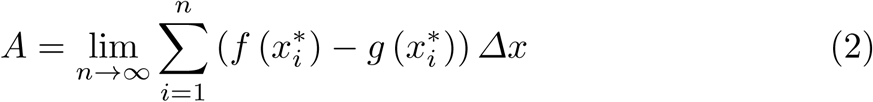

To demonstrate the shape of the internodes, five basal (the most developed) internodes were chosen. Volumes were measured as above and quadrilateral volumes were obtained, as below:

The area of a simple irregular quadrilateral (which approximates the inner area of a sugarcane internode) is determined according to Figure 6. Internally, the quadrilateral is divided by the diagonal q into an upper triangle *S*_1_ and a lower triangle *S*_2_.

**Fig. 6.**
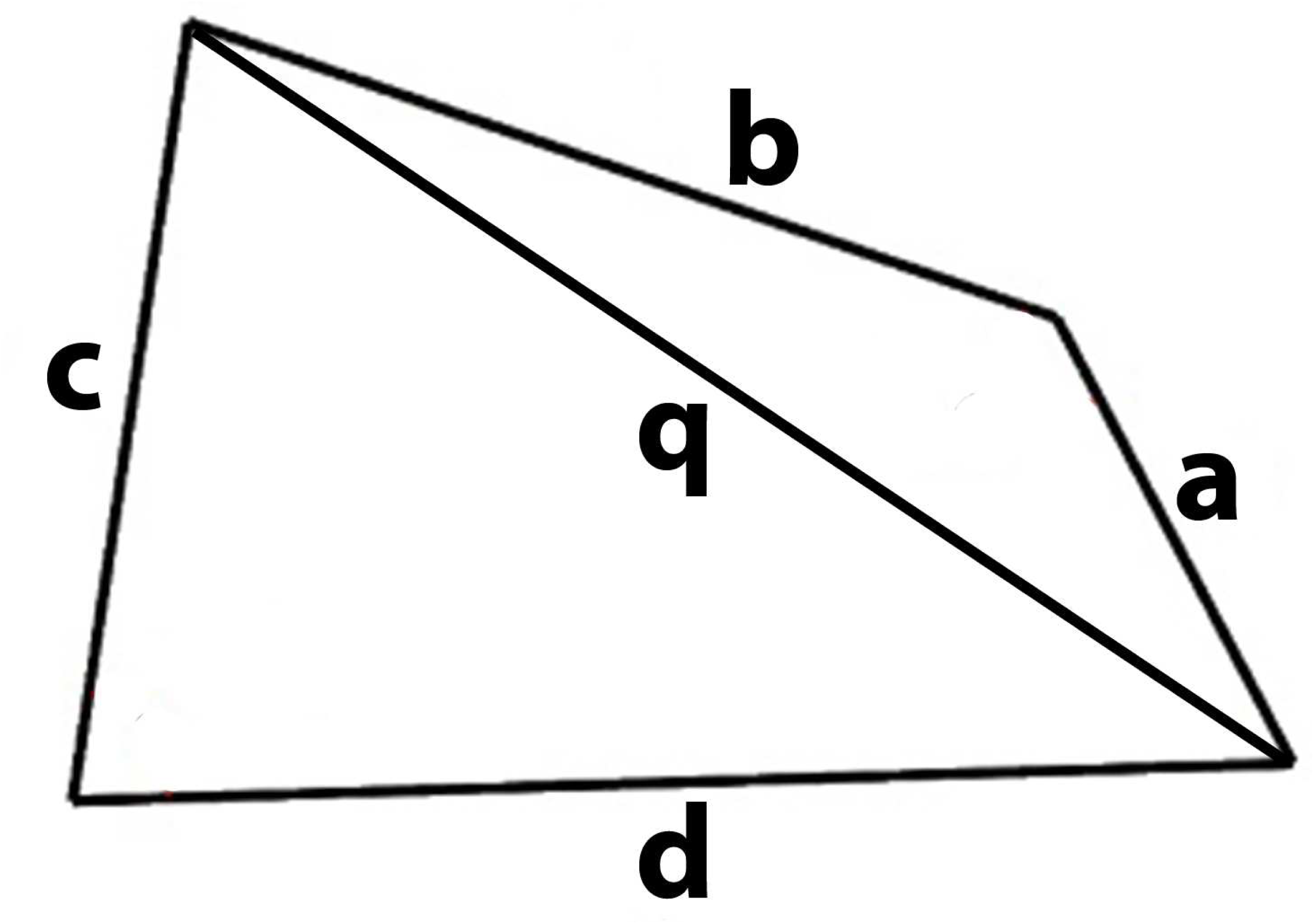
Image of a simple irregular quadrilateral to illustrate how its area is derived.

The semi-perimeters of these triangles are given by:

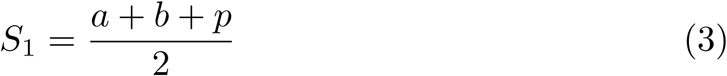

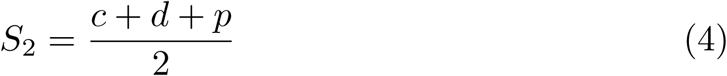

The area is then derived as:

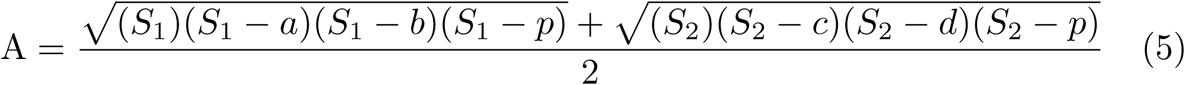

Correct internode volumes (from ICY and equation 2) were divided by simple quadrangle volumes, summed over five internodes and averaged. Where the number is >1 nodes are tumescent. Where the number ≈ 1 nodes are cylindrical, where the number < 1 nodes are bobbin-shaped.

### Plant Materials

Plant materials were obtained from the Mount Edgecombe campus of the South African Sugarcane Research Institute, Durban, a private collection on Gran Canaria, a distillery in Barbados and from searches of NCBI’s sequence reads archive (For full details see Supplementary Document 1). Mount Edgecombe materials were sequenced with 12 universal primers, and Illumina sequencing as described previously (Lloyd Evans et al. 2019). All plant materials were also sequenced using capture primers and MinION sequencing (Lloyd Evans and Hughes 2020). Full length 45s ribosomal RNA sequences were isolated from total RNA using capture primers prior to direct RNA sequencing with MinION (Lloyd Evans and Hughes 2020).

### R570 Sugarcane Centromere Enrichment and ITS Isolation

Based on the predicted sugarcane cenH3 sequence (Nagaki and Murata 2005) the peptide: MARTKHQAVRRPTQKPKKKLQKE was synthesised (University of Cambridge, Department of Biochemistry). This was provided to BioServe, Sheffield, UK for small scale monoclonal antibody synthesis. ChIP experiments were conducted according to published protocol (Nagaki et al. 2004). DNA was purified and fragmented prior to size fractionation at 10kbp (Storchevoi et al. 2020). Size fractionated DNA was further sub-selected using primers for sugarcane 45s ribosomal RNA sequences prior to magnetic bead extraction and MinION sequencing. MinION reads were further subset by mapping to sugarcane reference 45s ribosomal RNA sequences with minimap2 prior to assembly with Canu.

As the following methods have been published previously only a brief summary of the methods is given here. However, the full methods are available as Supplementary Document 2.

### Sequence Assembly

Illumina data were assembled using a bait and assemble methodology (Lloyd Evans and Joshi 2016) and MinION/PacBio data were assembled with Canu 2.0 (Bankevich 2012). Assembled chloroplasts and 45s ribosomal RNA sequences were combined with existing data (detailed in Supplementary Document 1) prior to alignment. Whole chloroplast sequences were aligned with SATE (Liu et al. 2009) whilst 45s rRNA regions were aligned with SATE and optimized with PRANK (Löytynoja et al. 2012), as described previously (Martin et al. 2017).

### Phylogenetics

Phylogenetic analyses for chloroplast and 45s ribosomal RNA regions were performed as described previously (Lloyd Evans and Hughes 2020) using IQ-Tree (Nguyen et al. 2015) and Mr Bayes (Huelsenbeck and Ronquist 2001). The same applications were employed to obtain branch supports. Final phylogenies were drawn with FigTree (http://tree.bio.ed.ac.uk/software/figtree/), and Adobe Illustrator.

## 6 List of abbreviations

creole, sugarcane, phylogenetics, history, Saccharum cultum

## 7 Availability of data and materials

All sequences generated in this study have been submitted to ENA under the project accessions PRJEB38889 (for chloroplast assemblies) and PRJEB38888 45s RNA assemblies. Sequence alignments and base phylogenies are available from (DOI). Computer code developed for the project is available from GitHub: https://github.com/gwydion1/bifo-scripts.git. Sequence alignments, third party assemblies and base phylogenies are available from Zenodo (DOI).

## 10 Authors’ contributions

DLlE conceived the experiment, performed the analysis and historical research, analyzed the data and wrote the paper. SVJ and DLlE secured the funding.

## 11 Acknowledgements

We would like to thank Mrs B Hughes for sourcing the Creole sugarcane samples from the Bahamas and Barbados.

