## Supplementary Document 1 for "Origins, History and Molecular Characterization of Creole Cane"

#### Ibn al Batar 14C

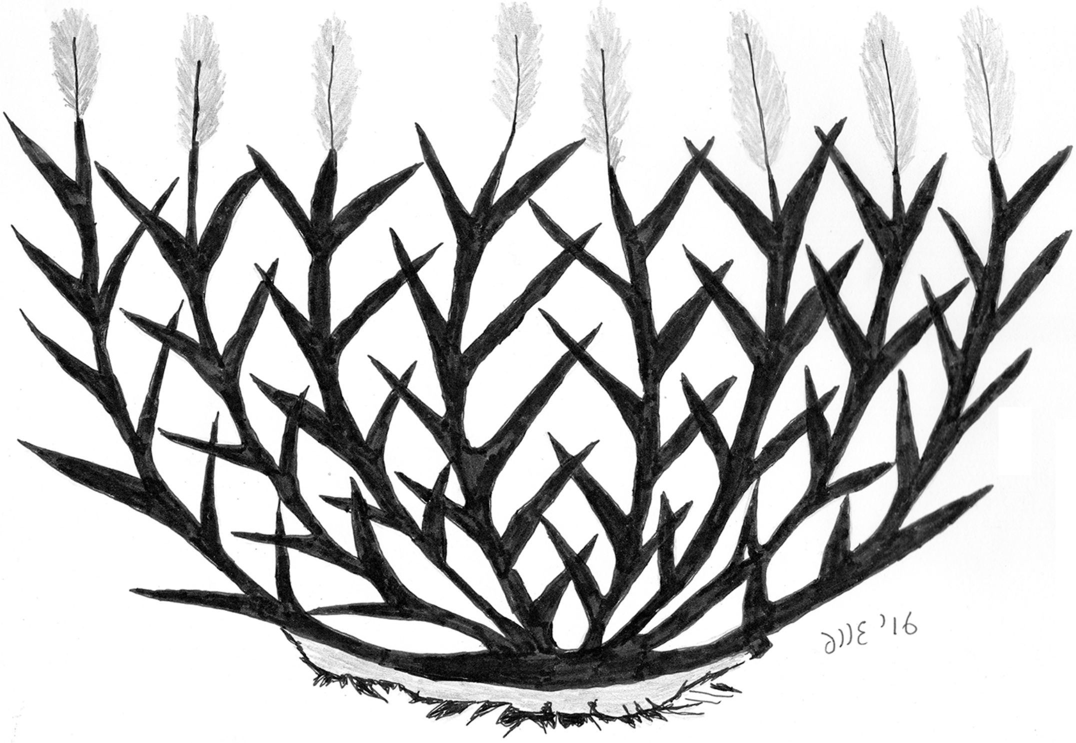

#### Tacuinum Sanitatis 14C

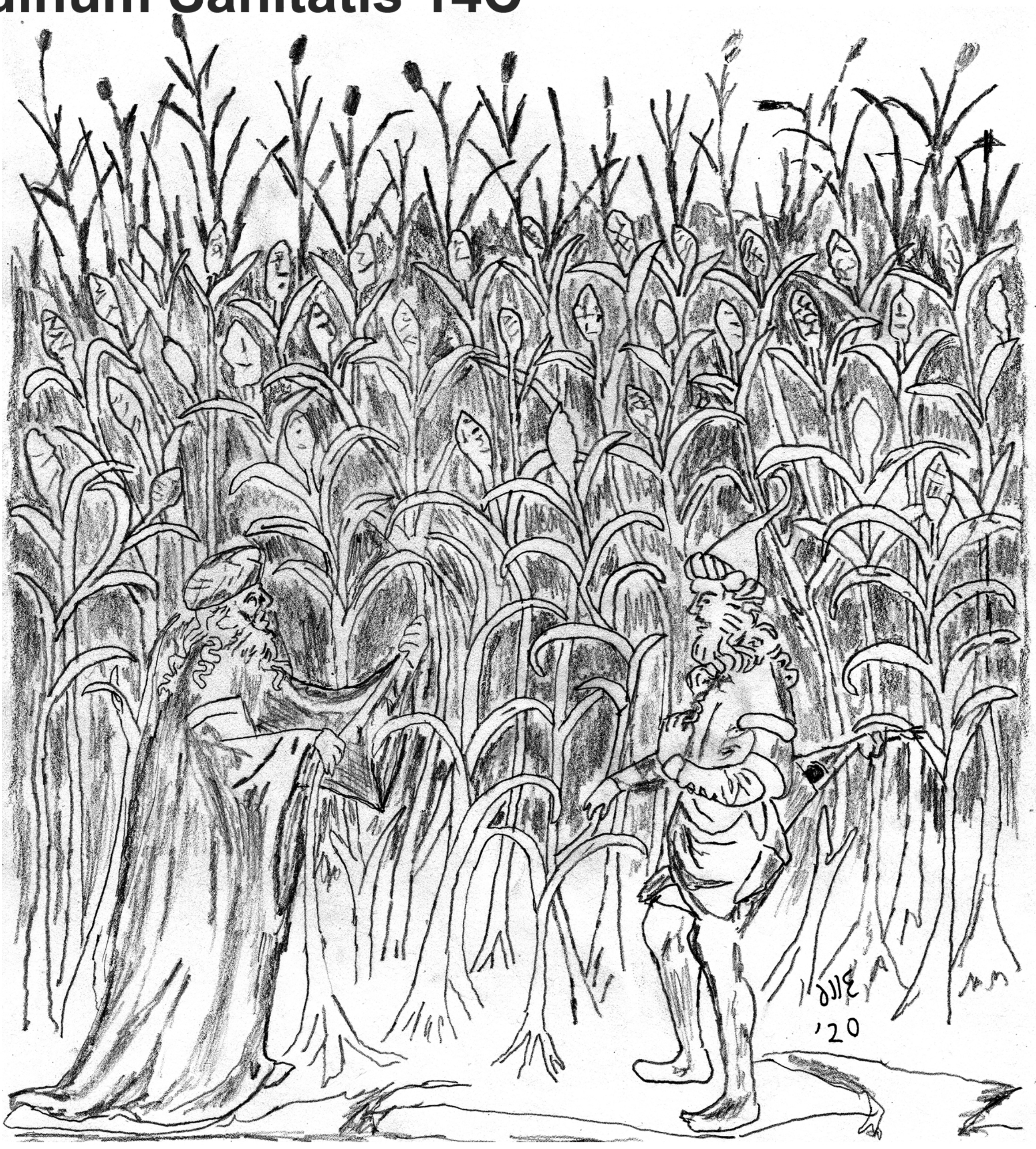

**Münster 1570**

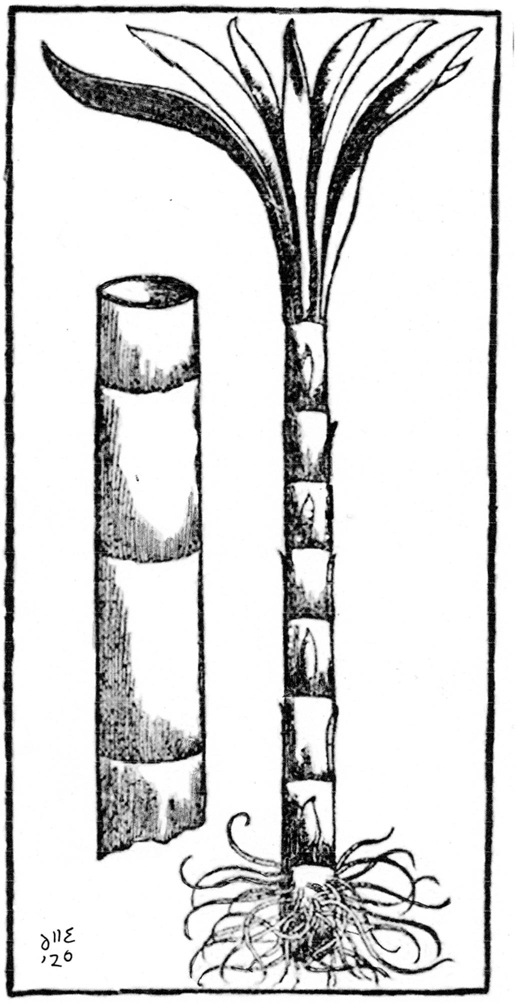

**Daléchamps 1587**

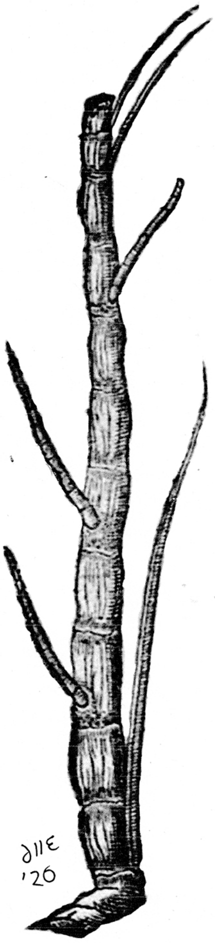

**Eckhout 1641**

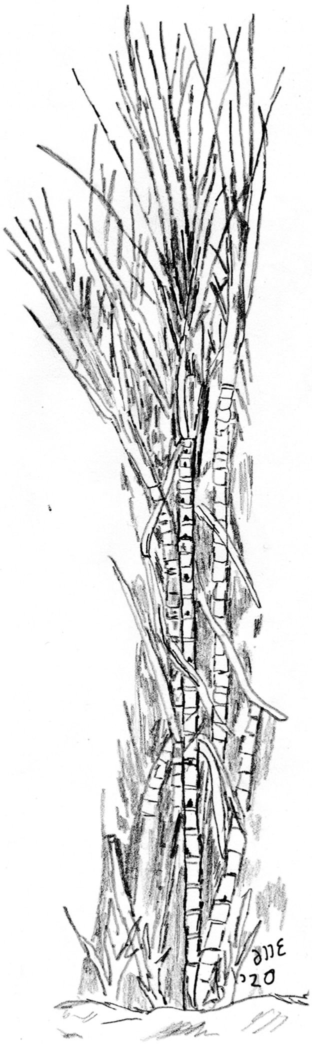

**Piso & Marcgrav 1648**

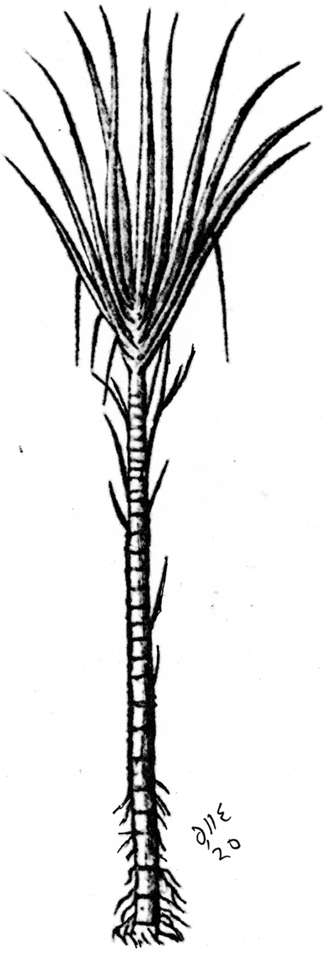

**Nieuhof 1682**

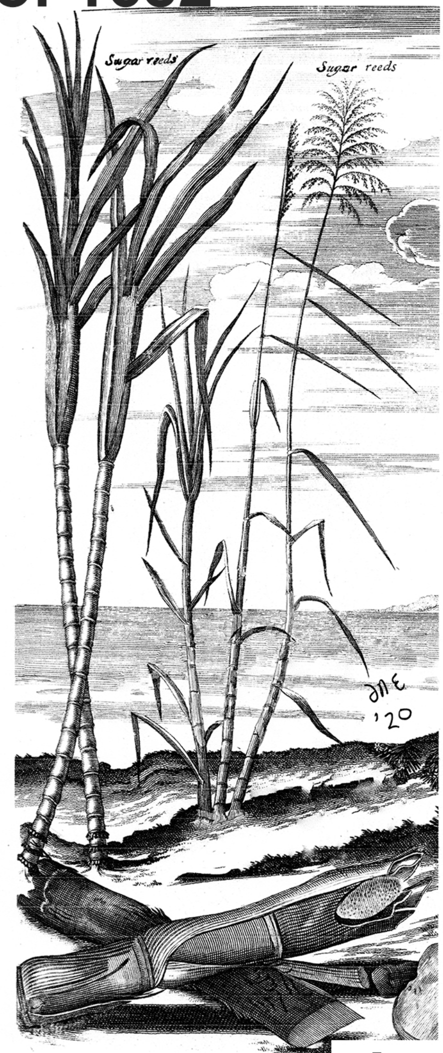


**Munting 1696**

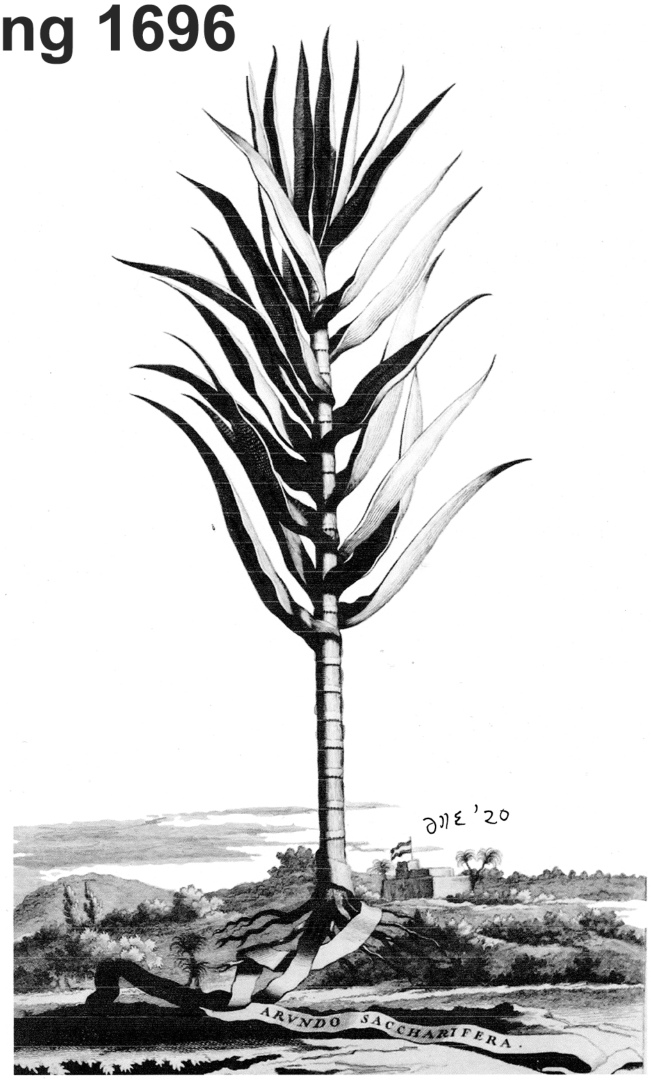


**Zwinger 1696**

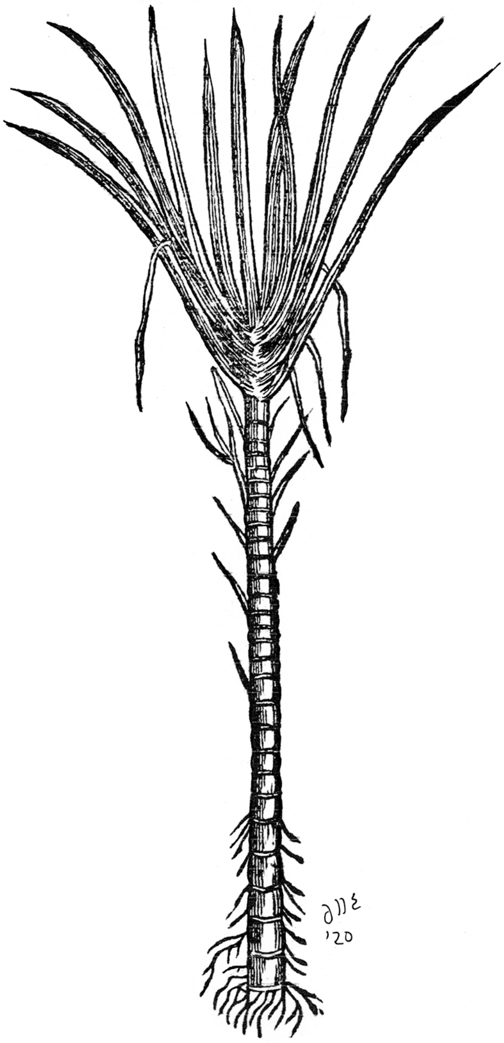

**de Quélus 1719**

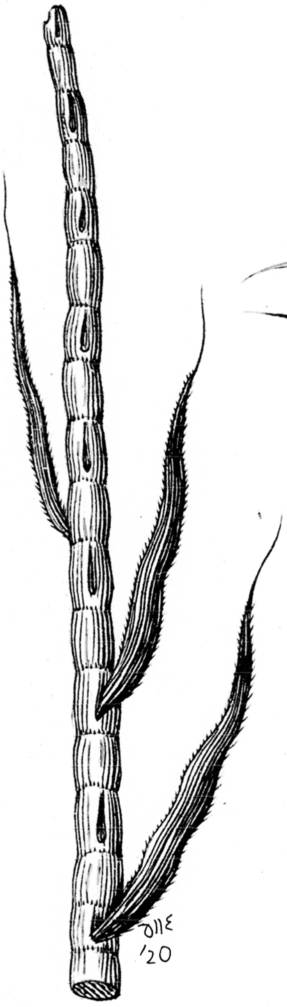

**Hughes 1750**

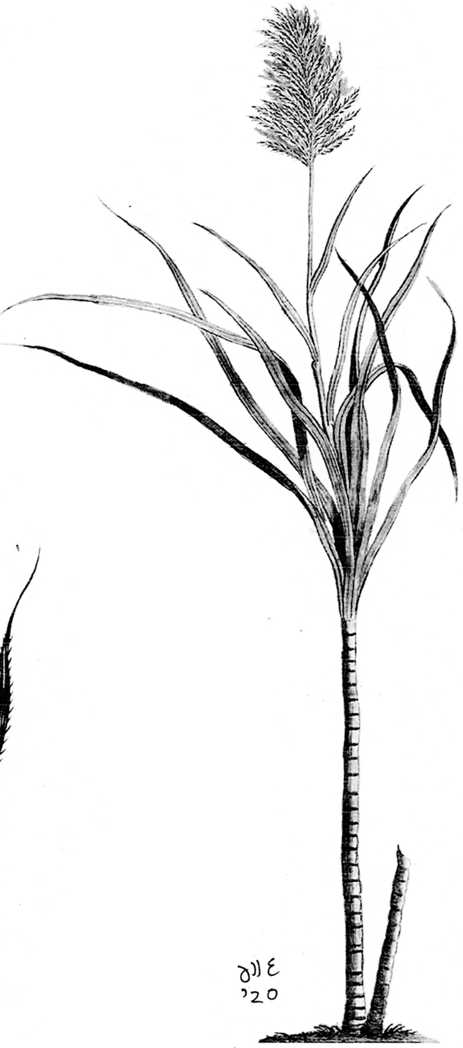

Herlein 1718

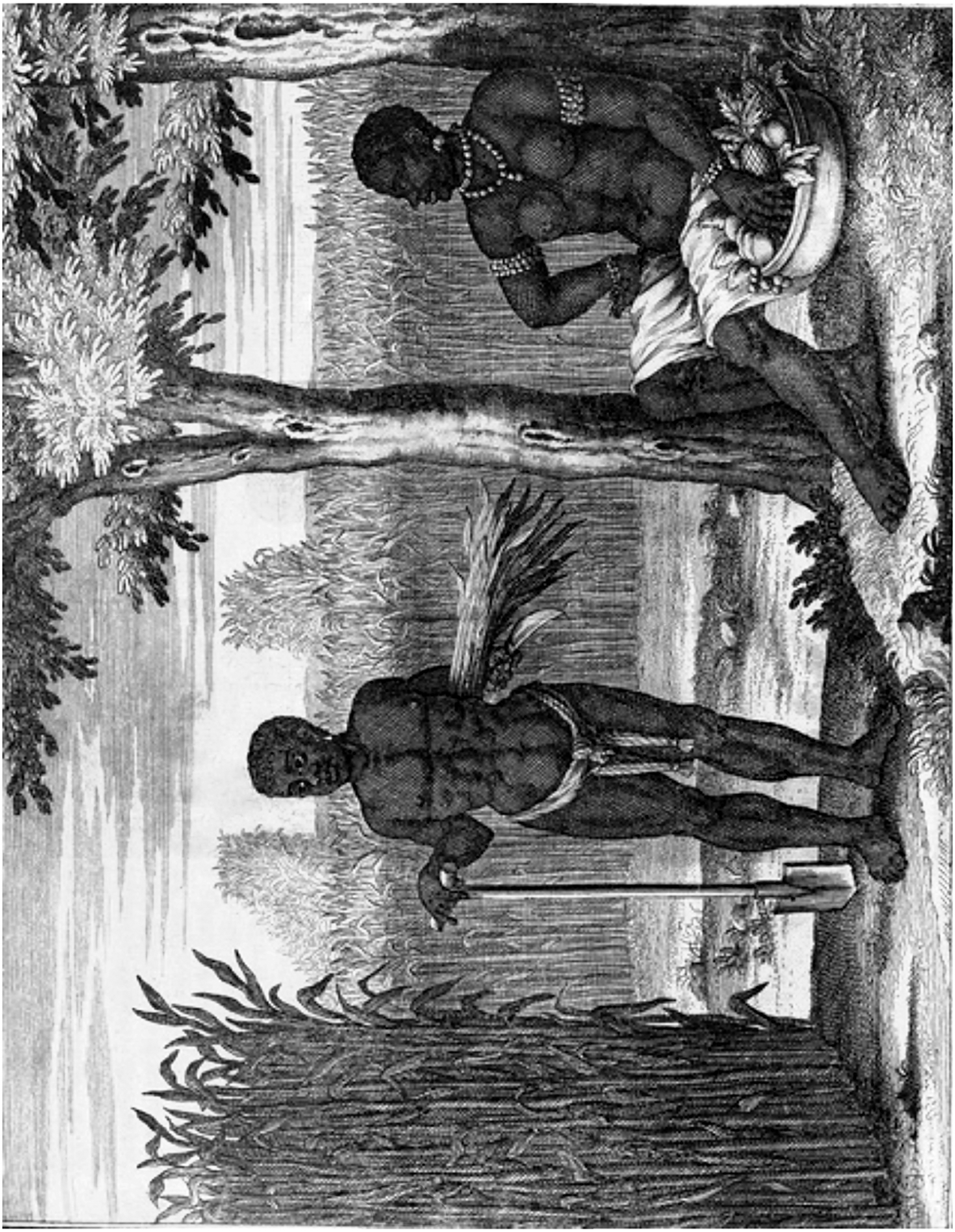

**Hinton 1749**

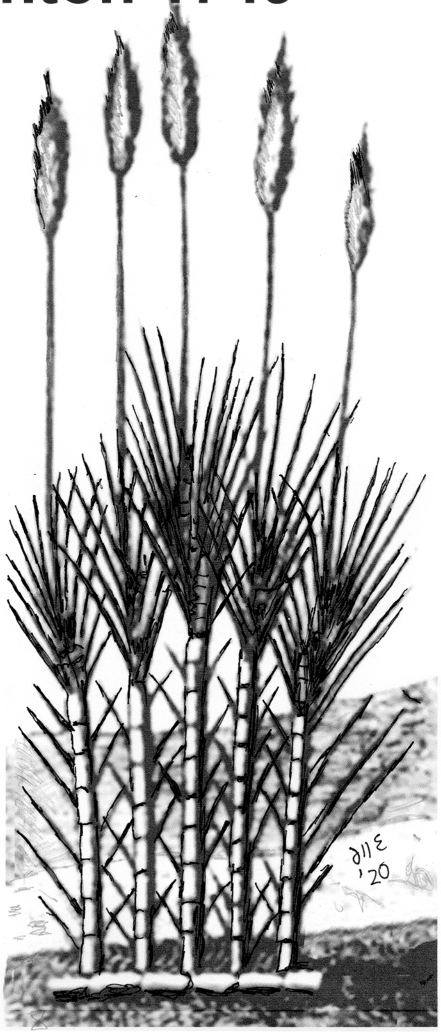

**Terrenise 1763**

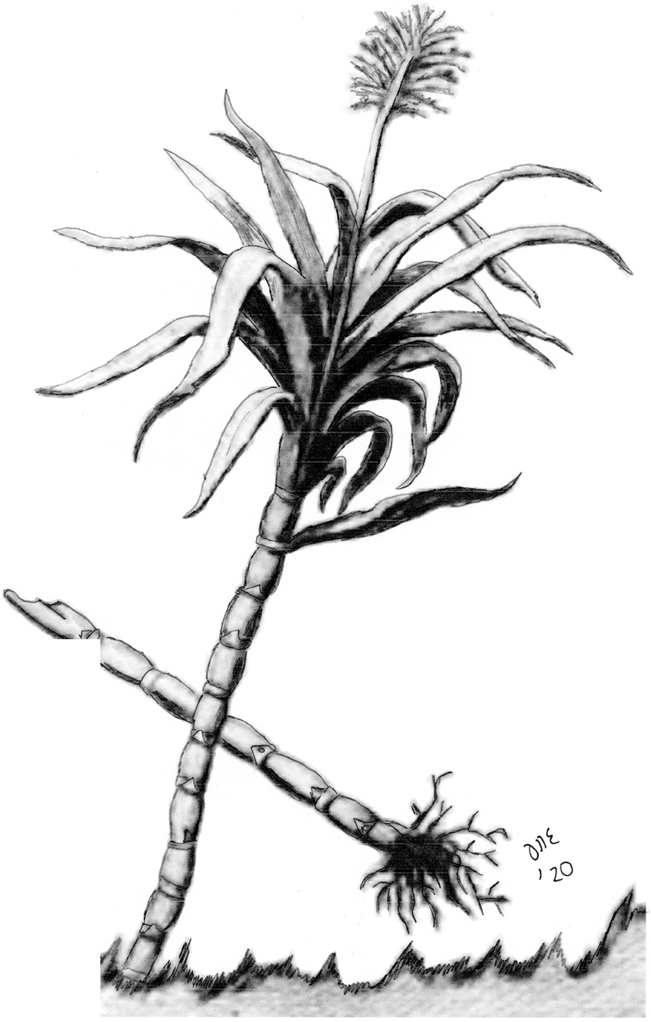

**Gersault 1768**

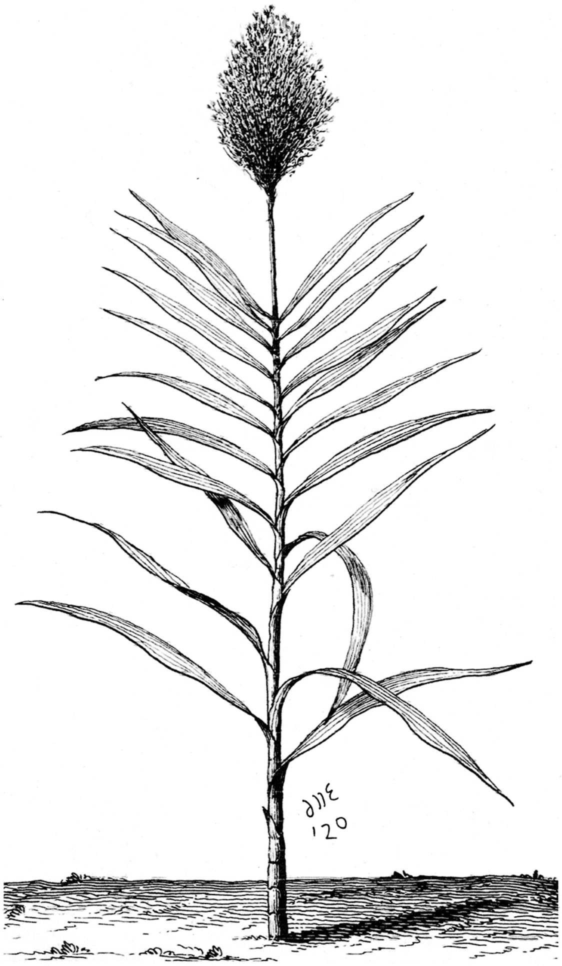

**de la Coture 1790**

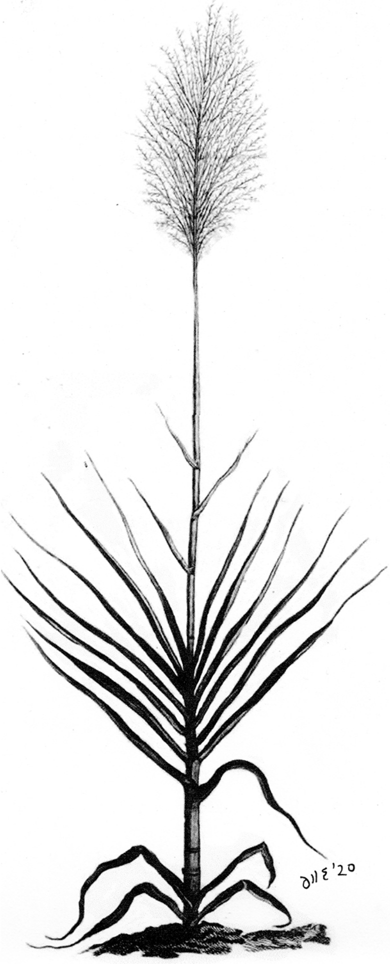

**Stedman 1791**

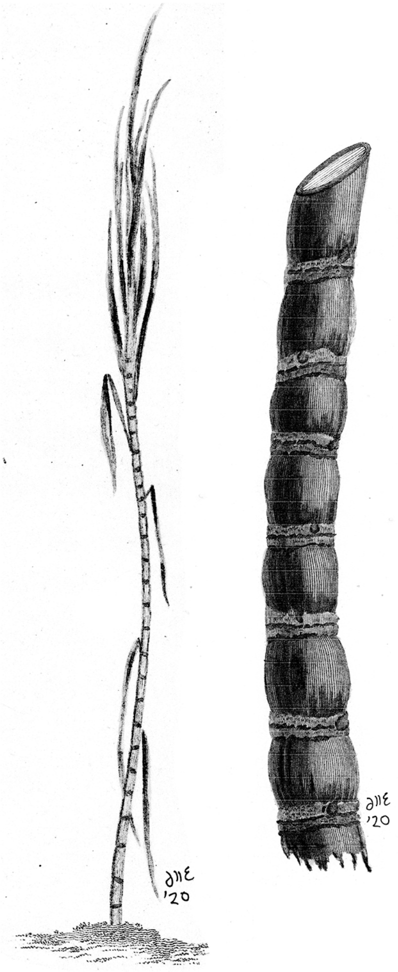

### Woodville 1790

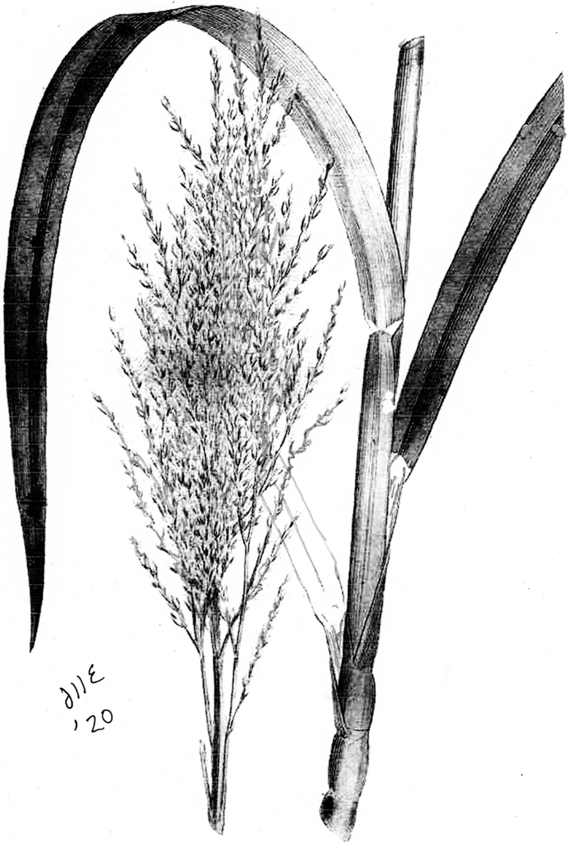

### Oxholm 1797

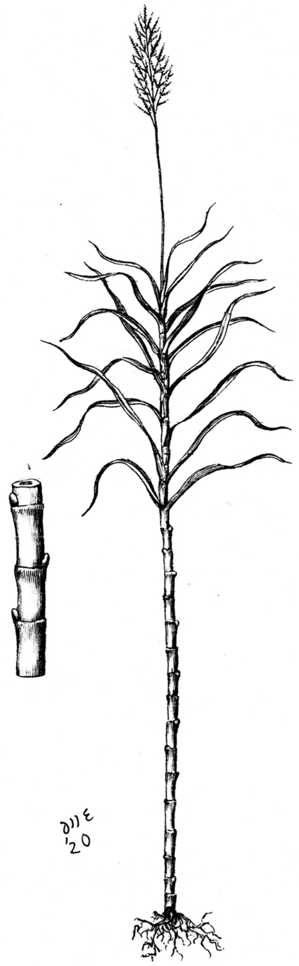

### Descourtilz 1829

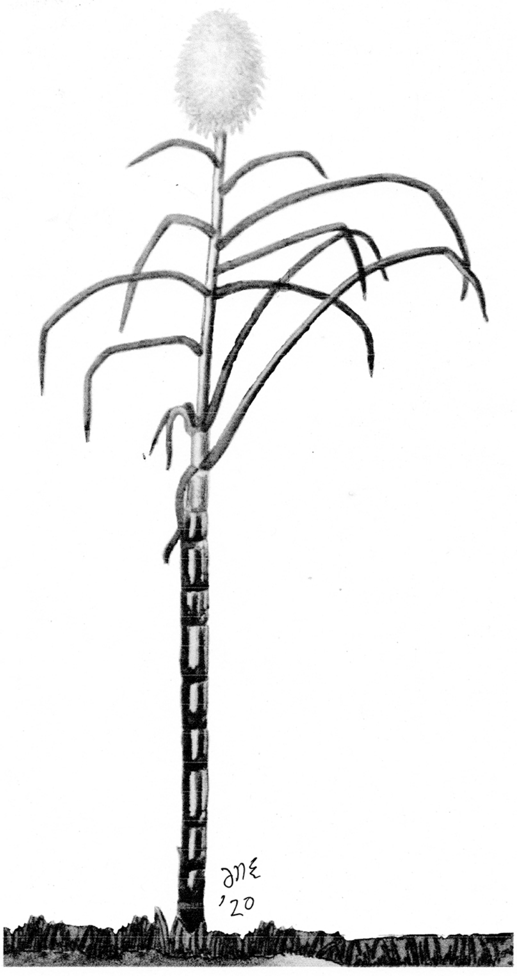

### Hooker 1830

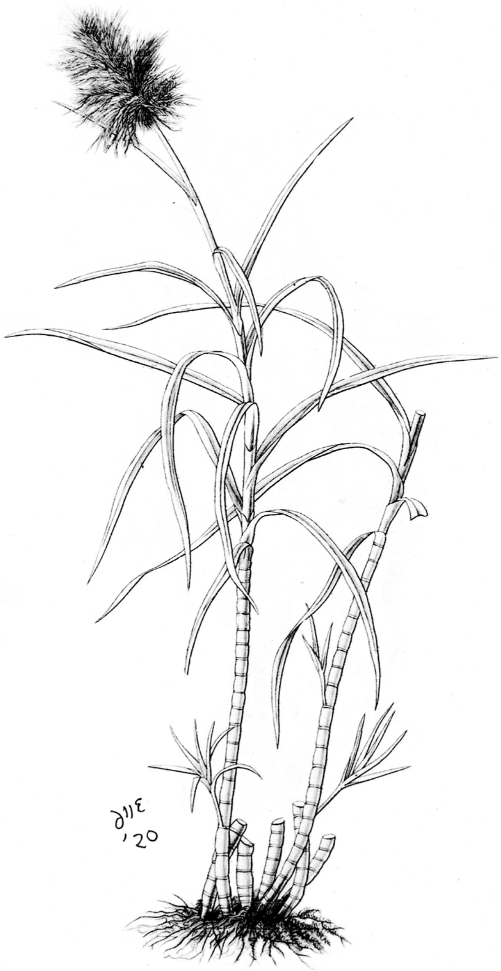

**Dietrich 1831**

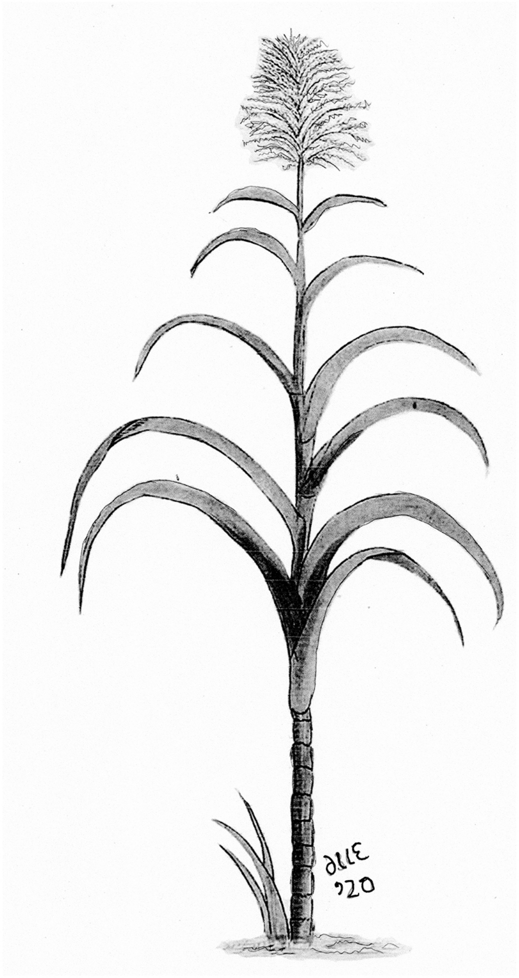

**Flschmann 1848**

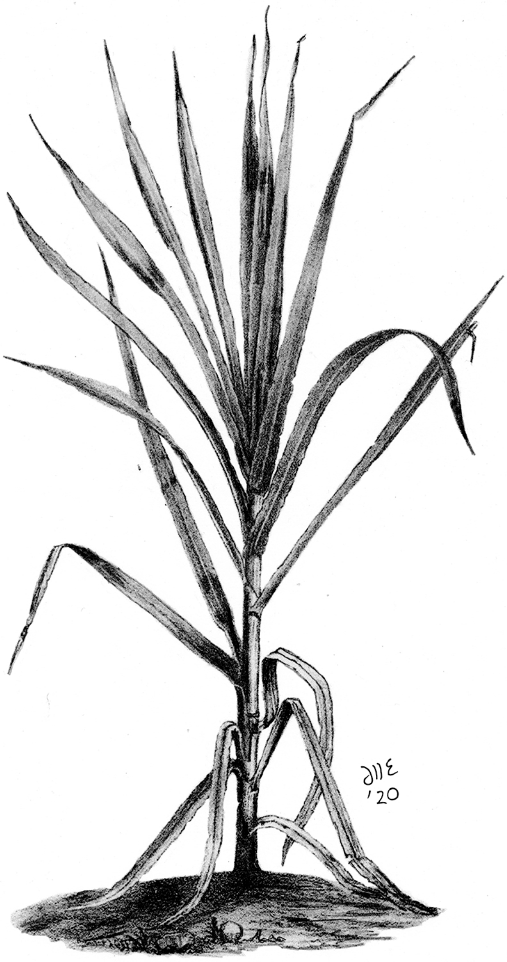

**Heck 1851**

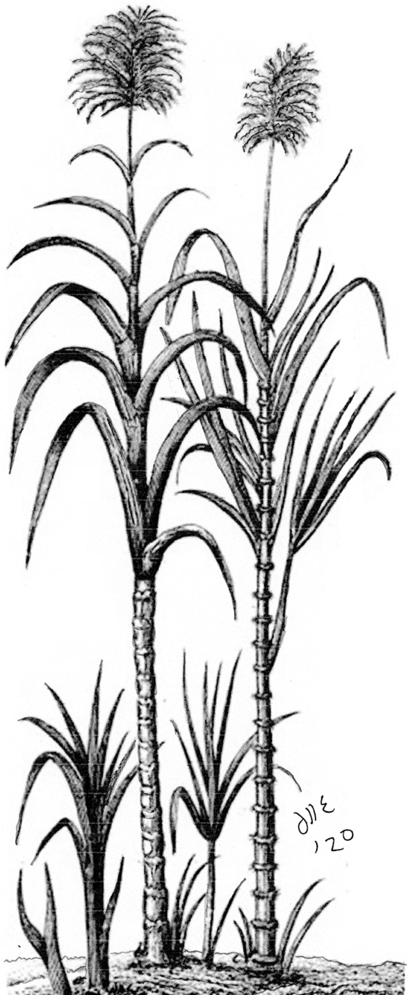

**Deerr 1921**

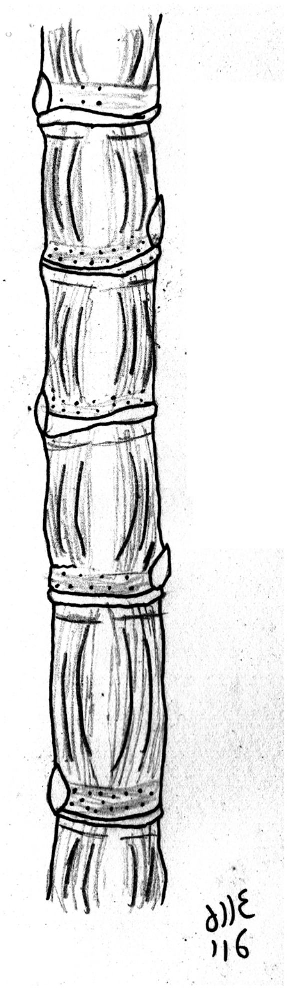

#### **Image Sources**:

Münster S. 1570. *Cosmographia Universalis*. Henry Durch. Basel.
Note: The First Edition of the Cosmographia was published in 1544 and though Sebastian Münster himself died in May 1552, his Cosmographia continued to be published until 1628. Sugarcane first appears in the 1570 edition, above a map of Malta. After Münster's death Heinrich Petri continued with the editions until his death in 1579.

Daléchamps J. 1587. *Historia Generalis Plantarum*. Lugdumi: Apud Gulielmum Rovilium.

Eckhout A. 1641. Mulatto Man, Oil on Canvas. National Museum of Denmark.

Piso W & Marcgrave G. 1648. *Historia naturalis Brasiliae: in qua non tantum plantæ et animalia, sed et indigenarum morbi, ingenia et mores describuntur et iconibus supra quingentas illustrantur*. Amsterdam : Lud Elzevirum

Nieuhof J. 1682. *Johan Nieuhofs Gedenkweerdige Brasiliaense zee- en lant-reize*. Amsterdam.

Munting A. 1696. *Naauwkeurige beschrijving der aardgewassen*. Vol I. Leyden: Pieter vander AA; Utrecht: Francois Halma.

Zwinger T. 1696. *Theatrum botanicum das ist: Neu vollkommenes Kräuter-Buch*. Basel.
Note, this new woodcut is based on the original image of Piso and Marcgrave, but as is common for a woodcut or engraving taken from an original, the image is printed reversed. The shape of the internodes are adjusted slightly, too.

Herlein JD. 1718. *Beschryvinge van de volk-plantinge Zuriname*. Leeuwarden: Meindert Injema (Figure 8, Plate 3).
Note: Though this image of a male and female slave has sugarcane on the side and in the background the images are not diagnostic as the plants are too tightly packed and nodes and internodes are not delineated. This is the earlest representation of sugarcane from the Dutch colonies. However, the image of sugarcane itself is not diagnostic ans nodes and internodes are not delineated.

This image also requires some comment from an art historical perspective. The figure on the left, though now depicted as male was probably originally female. Note the width of the hips, the pinched waist and the feminine features. The figure on the right, though female now might have been male originally (note the masculine head). It simply would not have been done to show the woman working. This image also presages future Dutch depictions of slavery. Note the happy slaves --- no indication here of the truly appalling conditions of slaves in Dutch plantations. Also note the absence of slave brands, which would have been ubiquitous. This is a sanitized version of the truth meant for consumption at home. The slaves were happy with their lot, they were treated well and lived in an idyllic setting. The Dutch people could be happy that the sugar welath coming to their nation did not originate from maltreatment of the slaves. This is also a rosy and sanitized image of slavery that we also see in the artwork of Eckhout (c. 1740s).

de Quélus [A]. 1719. *Histoire Naturelle du cacao et du sucre*. Paris: Laurent d'Houry

Anon. 1749. Image titled: A representation of the sugar-cane and the art of making sugar.. The universal magazine of knowledge and pleasure .... London : Published ... according to Act of Parliament, for John Hinton, 1749.

Hughes G. 1750. *The Natural History of Barbados.* London

Terrenise GM. 1763. "Canna da zucchero/Plantazione di zucchero," engraving. In *Il gazzettiere americano contenente un distinto ragguaglio di tutte le parti del Nuovo Mondo*. Livorno.

de Gersault FAP, Geoffroy E-F. 1767. *Description, vertus et usages de sept cents dix-neuf plantes, tant étrangères que de nos climats.* Vol I. Paris :P.F. Didot le jeune

de la Coture J-F D. 1790. *Précis sur la canne et sur les moyens d'en extraire le Sel essentiel, suivi* *de plusieurs mémoires sure le Sucre, sur le Vin de Canne, sur l'Indigo, sur les habitations & sur l'état actuel de Saint-Domingue*. Paris: Duplain

Woodville W. 1790. Medical Botany. London: James Phillips.

Stedman JG. 1791. Narrative, of a five years' expedition, against the revolted Negroes of Surinam ... Vol. I. J. Johnson, St. Paul's Church Yard. Image plate 34, entitled: Four Stages of Sugar Cane Growth.

**Beyond the 1790s Tahitian, Batavian (Javanese) cane had reached the New World, thus the followiing images, though the majority depict Creole cane should be treated with some circumspection.**

Oxholm PL. 1797. *De Danske Vestindiske öers tilstand i henseende til population, cultur og finance-forfatning, i anledning af nogle breve fra St. Croix.* Kiöbenhavn: Trykt hos directeur Johan Frederik Schultz. fold-out plate II; following p. 84
Note: This is not Creole cane. The sparser infloresence and bobbin shaped internodes indicates that it is probably ribbon cane (Guingham).

Descourtilz ME. *Flore [pittoresque et] médicale des Antilles*. V 7. Paris: Pichard
Note: This is not Creole cane, but Ribbon Cane (Guingham)

Hooker WJ. 1830. Botanical Miscellany. J Murray. London

Dietrich DNF. 1831 *Flora Medica*. Jena: August Schmid

Fleischmann CL. 1848. Report on Sugar Cane and its Culture. In: Annual Report (Agriculture) U.S. Commisioner of Patents for 1848. Washington: G.P.O., pp 274-338.

Heck JG. 1851. *conographic Encyclopedia of Science, Literature and Art*. Translated from the German, with additions, and edited by Spencer F. Baird. 4 Bände. Garrigue, New York

Deerr N. 1921. Cane Sugar. London: Norman Roger

*All images ©Dyfed Lloyd Evans, reproduced with permission*
